# LRRK2 mediates tubulation and vesicle sorting from membrane damaged lysosomes

**DOI:** 10.1101/2020.01.23.917252

**Authors:** Luis Bonet-Ponce, Alexandra Beilina, Chad D. Williamson, Eric Lindberg, Jillian H. Kluss, Sara Saez-Atienzar, Natalie Landeck, Ravindran Kumaran, Adamantios Mamais, Christopher K. E. Bleck, Yan Li, Mark R. Cookson

## Abstract

Mutations in the leucine rich repeat kinase 2 (*LRRK2*) gene are a cause of familial and sporadic Parkinson’s disease (PD). Nonetheless, the biological functions of LRRK2 remain incompletely understood. Here, we observed that LRRK2 is recruited to lysosomes that have a ruptured membrane. Using unbiased proteomics, we observed that LRRK2 is able to recruit the motor adaptor protein JIP4 to permeabilized lysosomes in a kinase-dependent manner through the phosphorylation of RAB35 and RAB10. Super-resolution live cell imaging microscopy and FIB-SEM revealed that once at the lysosomal membrane, JIP4 promotes the formation of LAMP1-negative lysosomal tubules that release membranous content from ruptured lysosomes. Released vesicular structures are able to interact with other lysosomes. Thus, we described a new process that uses lysosomal tubulation to release vesicular structures from permeabilized lysosomes. LRRK2 orchestrates this process that we name LYTL (LYsosomal Tubulation/sorting driven by LRRK2) that, given the central role of the lysosome in PD, is likely to be disease relevant.

## INTRODUCTION

Mutations in *LRRK2* are a relatively common cause of familial late-onset Parkinson’s disease (PD)(*1*),(*2*), and variations at the *LRRK2* locus have also been linked to the more numerous sporadic PD(*3*),(*4*),(*5*). *LRRK2* encodes Leucine-rich repeat kinase 2, a large protein with both GTPase and kinase activity. The majority of proven pathogenic mutations are located in the ROC-COR bidomain that controls GTP hydrolysis and kinase activity and the majority of evidence suggests that mutations lead to a toxic function of the protein(*6*),(*7*). At the cellular level, evidence suggests that LRRK2 can regulate membrane trafficking events via phosphorylation of a subset of RAB GTPases(*8*),(*9*),(*10*), although the precise relationship(s) between LRRK2 mutations, RAB phosphorylation and neurodegeneration remain uncertain.

Recent genetic data have pointed to the lysosome as a crucial organelle in PD, as mutations in genes encoding for lysosomal proteins have been identified in familial cases of PD(*11*) and have been nominated as a risk factor for sporadic PD(*4*), leading to the suggestion that PD should be considered a lysosomal disease(*12*). An accumulation of lysosomal damage with age in kidneys, which normally express high levels of LRRK2, has been documented in knockout mice(*13*),(*14*),(*15*). Pathogenic LRRK2 mutations affect lysosomal structure and function in cultured astrocytes(*16*) and other cell types(*17*),(*18*),(*19*),(*20*). However, the mechanistic basis by which LRRK2 affects lysosome function is unclear. Additionally, because LRRK2 can be localized to a wide range of other membrane-bound structures in cells(*8*),(*21, 22*), whether LRRK2-mediated lysosomal defects are primary events or secondary effects driven by toxicity to other cellular components is uncertain.

Here, we describe for the first time that LRRK2 translocates to the membrane of permeabilized lysosomes, leading to the phosphorylation and recruitment of RAB35 and RAB10. As a consequence, both RAB proteins promote the translocation of the motor adaptor protein JIP4. JIP4 is present in, and helps to form, LAMP1-negative tubular structures stemming from lysosomes. Live cell super-resolution microscopy reveals that these tubules bud, extend and release small vesicular structures, suggesting a scenario where ruptured lysosomes sort membranous content that can then interact with other lysosomes. We call this newly described process LYsosomal Tubulation/sorting driven by LRRK2 (LYTL).

## RESULTS

### LRRK2, along with the motor adaptor protein JIP4, gets recruited to the membrane of a subset of lysosomes

To understand how LRRK2 might affect lysosomal function, we first examined the localization of LRRK2 in cells. For these experiments, we decided to use mouse primary astrocytes as (i) primary astrocytes express LRRK2 endogenously (unpublished data from our lab, Fig. S1A), (ii) their flattened morphology allows us to monitor membrane trafficking events and (iii) they have been proposed to play a role in the pathobiology of PD through non-cell autonomous effects on microglia and neurons(*23*). Exogenously expressed LRRK2 has been widely reported in the literature to have a diffuse cytosolic distribution(*24*),(*25*),(*8*), however when expressed in mouse primary astrocytes, LRRK2 colocalized with the late-endosomal/lysosomal membrane marker LAMP1 in about half of the cells examined. Specifically, LRRK2 is present at a subset of late-endosomes/lysosomes (LE/LYS) (2.723 ± 0.408 LRRK2-positive LE/LYS structures per cell) (Fig.1A) that are also positive for the late-endosomal/lysosomal marker RAB7 and the lysosomal marker LAMP2. These structures only occasionally contain the lysosomal enzyme cathepsin B (CTSB), while being negative for the early-endosomal marker EEA1 (Fig. 1B). The combination of the presence of several LE/LYS markers with variable levels of CTSB suggests that LRRK2 is recruited to LYS with a low degradative capacity(*26*).

**Figure 1.**
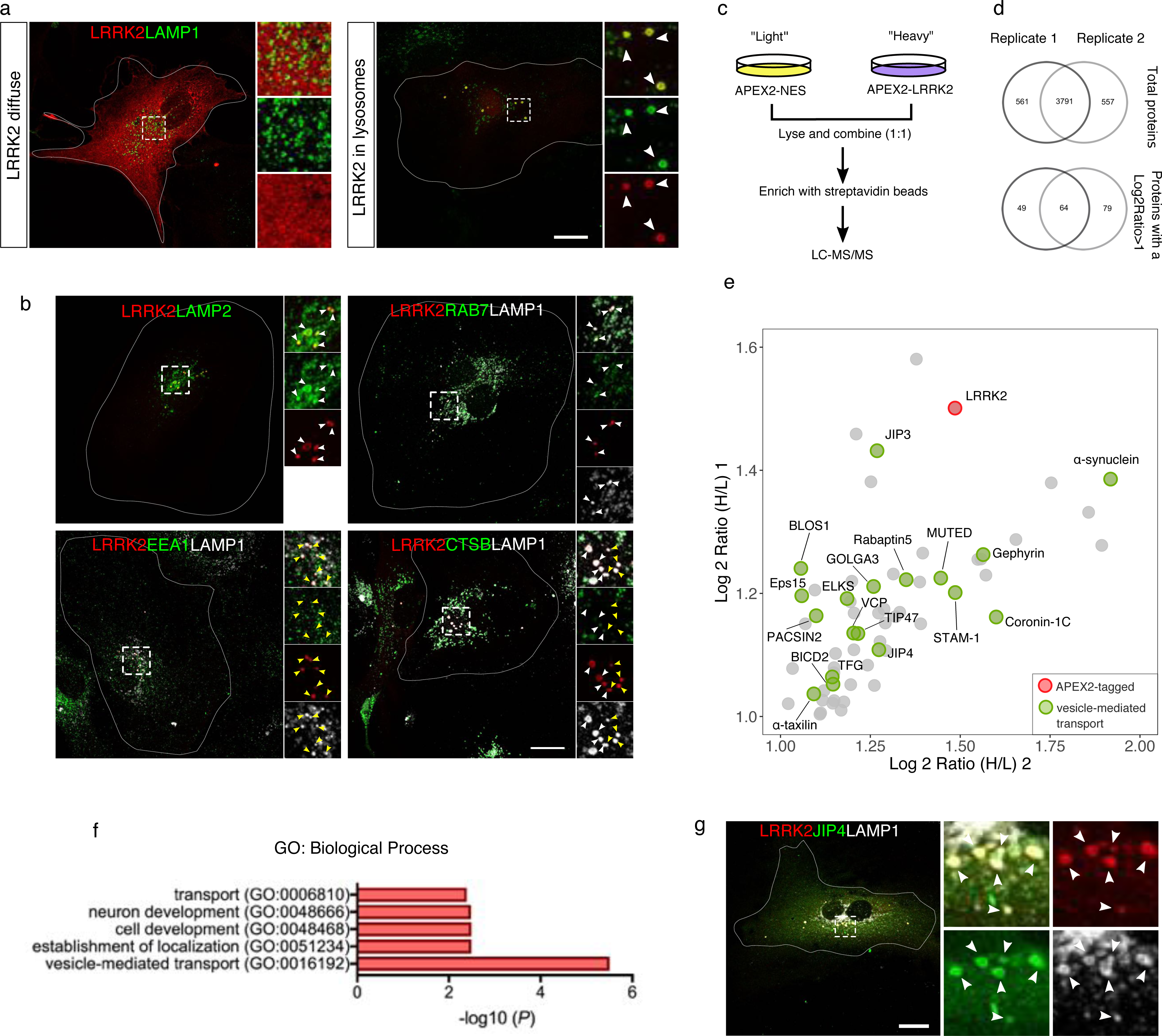
LRRK2 and JIP4 are localized in the membrane of a subset of lysosomes in primary astrocytes. **A**. Representative confocal images of 3xflag-LRRK2 (red) and LAMP1 (green) expression in mouse primary astrocytes. **B**. Representative confocal images of astrocytes expressing 3xflag-LRRK2 (red) co-stained with LAMP2, RAB7, EEA1 and CTSB (green). **C**. Outline of the APEX2 proteomic approach to detect LRRK2-membrane interactors. HEK293FT cells were transfected with 3xflag-APEX2-LRRK2 (heavy amino acids SILAC media) and with 3xflag-APEX2-NES (light amino acids SILAC media). **D**. Venn diagram showing the number of common proteins detected in both replicates and the number of proteins selected as “candidates” in both replicates. **E**. Scatter plot depicting the 64 LRRK2-interacting candidates from the APEX2 screening. LRRK2 is marked in red, and proteins involved in vesicle-mediated transport are marked in green. **F**. Gene Ontology search of the top 5 enriched terms for Biological Process of the 64 LRRK2 potential interacting partners, with *p.*values subjected to Bonferroni correction. **G**. Representative confocal image of an astrocyte expressing 3xflag-LRRK2 (red), GFP-JIP4 (green) and stained for LAMP1 (white). White arrowheads show colocalization and yellow arrowheads show structures without localization. Scale bar= 20 µm.

We next questioned whether LRRK2 enzymatic activity could play a role in its recruitment to the LYS by expressing the hyperactive and pathogenic mutations G2019S (kinase domain) and R1441C (GTPase domain), along with the artificial inactive mutations K1906M (kinase domain) and T1348N (GTPase domain) (Fig. S1B). An increase in LRRK2-positive LYS per cell was observed in both hyperactive mutations compared to the wild-type (WT) form, as well as a strong reduction in both inactive mutations (Fig. S1C). Domain specific LRRK2 constructs (ΔHEAT, ΔWD40, HEAT, ROC-COR-Kinase, ROC-COR) (Fig. S1D) showed much less recruitment to LYS compared to the full-length construct (Fig. S1E, F). However, comparing the domains to each other, we detected a higher amount of lysosomal LRRK2 in the HEAT and ΔWD40 domains compared to the other three domains (Fig. S1E,G), suggesting that although every domain seems to be important to maintain LRRK2 at the lysosomal membrane, its recruitment likely happens through the N-terminal Armadillo domain.

These results suggest that while not all LRRK2 is lysosomal, it is likely to play a role at a subset of these organelles. To identify lysosome-specific functional interactors of LRRK2, we used the unbiased proximity-based biotinylation protein-protein interaction method ascorbate peroxidase 2 (APEX2)(*27*), followed by quantitative proteomics in cells with stable isotope labeled amino acids (SILAC). As a cytosolic control to exclude non-specific interactions, APEX2 was tagged to a nuclear export signal (NES) (Fig. 1C). Both vectors (APEX2-3xFLAG-LRRK2 and APEX2-3xFLAG-NES) were successfully validated using immunostaining and western-blot (Fig. S2A, B). Out of the mass-spec hits found in two independent replicates with the LRRK2 vector but not APEX-NES, sixty-four proteins were selected as possible candidates for LRRK2 interaction (Fig. 1D, E, Fig. S2C). Interestingly, among these candidates, six have already been linked to LRRK2 in previous studies (ɑ-synuclein, JIP3, JIP4, Coronin-1C, Gephyrin and PACSIN2)(*15*),(*28*),(*29*), suggesting that this method can detect authentic LRRK2 interactors. Fifteen of the candidate proteins have a role in vesicle-mediated transport (Fig. 1F) and six have been linked to lysosome biogenesis or dynamics in previous studies (BLOS1, Muted, JIP3, JIP4, BICD2, and STAM1). To further prioritize within the lysosomal-related hits, we tested whether they were recruited to LRRK2-positive LYS in astrocytes. JIP4 was the only candidate that colocalized with LRRK2-positive LYS (Fig. S2D, Fig. 1G). We were able to validate that there was a physical interaction between these two proteins by overexpressing LRRK2 and blotting for endogenous JIP4 in cells (Fig. S2E). We therefore considered JIP4, previously nominated by several independent laboratories using different techniques, to be a reliable LRRK2 interacting protein and a candidate for mediating any functional effects of LRRK2 on the lysosome.

### LRRK2 recruitment to lysosomes occurs as a response to lysosomal membrane rupture, independent of lysophagy

JIP4 (C-Jun-amino-terminal kinase-interacting protein 4) is a cytosolic scaffolding protein, associated with multiple aspects of vesicle-mediated transport by acting as an adaptor for both dynein and kinesin motor proteins(*30*). JIP4 has also been linked to stress response(*31*), we speculated that the LRRK2:JIP4 complex might respond to lysosomal damage. Such a role would be consistent with data described above showing that LRRK2-positive LYS have low levels of CTSB (Fig. 1B), which is seen when lysosomal membrane is ruptured and its contents leak into the cytosol(*32*). To test the hypothesis that lysosomal membrane damage may trigger LRRK2:JIP4 recruitment to lysosomes, we treated primary astrocytes with the strong lysosomotropic reagent L-leucyl-L-leucine methyl ester (LLOME). LLOME enters the cell via endocytosis and is transported to the lysosomes where it undergoes condensation by Cathepsin C, leading to lysosomal membrane rupture(*33*). Exposure of cells to 1 mM LLOME triggered a striking and time-dependent increase in LRRK2 recruitment to the lysosomal membrane (Fig. 2A). We confirmed that LRRK2 is recruited to ruptured lysosomes by showing that these structures are negative for both LysoTracker and Magic-Red CTSB (Fig. 2B, C), fluorescent probes that measure lysosomal pH and activity respectively.

**Figure 2.**
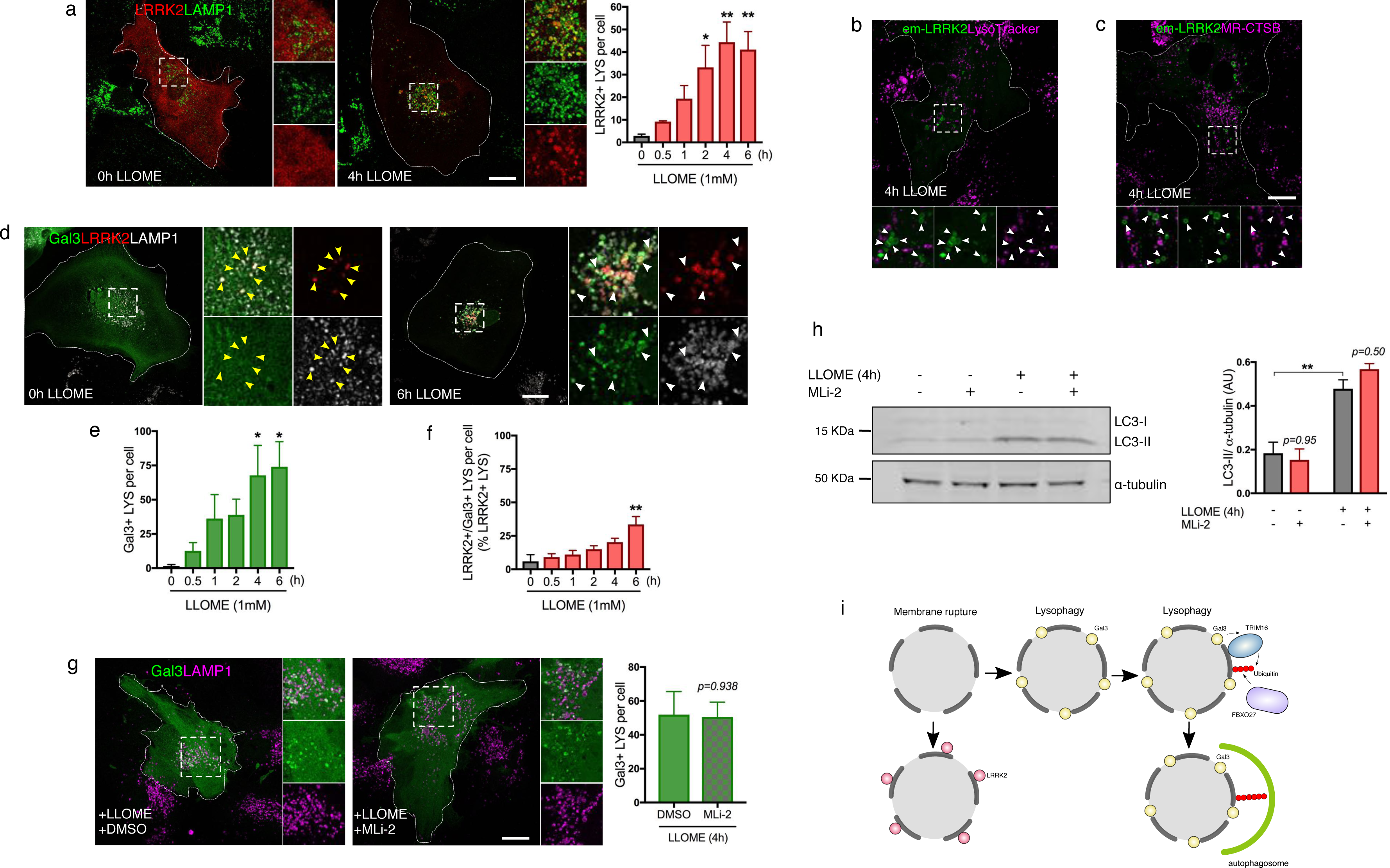
Lysosomal membrane permeabilization enhances LRRK2 recruitment to the lysosomal membrane, unrelated to lysophagy. A. Representative confocal images of astrocytes either untreated or treated with LLOME expressing 3xflag-LRRK2 (red) and LAMP1 (green). The histogram shows that the number of LRRK2-positive lysosomes per cell increases significantly after LLOME addition, compared to the untreated group. Data are mean ± SEM from 5 independent experiments with *n*= 20 cells per condition analyzed by one-way ANOVA (F (5,18)= 6.501, *p*= 0.0013). Multiple comparisons were evaluated using Dunnett’s *post-hoc* test. B-C. Representative live cell confocal images of astrocytes expressing Emerald-GFP-LRRK2 (green) exposed to LysoTracker Red DND-99 (magenta) (B) or Magic-Red CTSB (magenta) (C) and treated with LLOME for 4 h. White arrowheads show absence of colocalization between LRRK2 and the two lysosomal dyes. D-F. Representative confocal images of astrocytes expressing 3xflag-LRRK2 (red), EGFP-Gal3 (green) and LAMP1 (white) treated or not with LLOME (D). Only co-transfected cells were analyzed, yellow arrowheads indicate absence of colocalization while white arrowheads show colocalization E. Histogram shows the number of Gal-positive lysosomes per cell. Data are means ± SEM. One-way ANOVA with Dunnett’s post hoc for multiple comparisons was used, F (5,12)= 3.869, *p*= 0.0255, *n*= 12 cells per condition in 3 independent replicates. F. For colocalization analysis, 10-12 cells per condition were analyzed (with at least 3 LRRK2-positive lysosomes in each cell). The percentage of LRRK2-positive/Gal-positive lysosomes normalized by the total number of LRRK2-positive lysosomes was measured in each cell (in 3 independent replicates). Data are means ± SEM. One-way ANOVA with Dunnett’s test for multiple comparisons was used, F (5,12)= 6.605, *p*= 0.0036. G. Representative confocal images of astrocytes expressing EGFP-Gal3 (green) and LAMP1 (magenta), pre-treated with DMSO or MLi-2 and then incubated with LLOME. The number of Gal3-positive lysosomes per cell was measured. Data are means ± SEM. Unpaired *t-test* was applied (*p*= 0.938), *n*= 20-39 cells were analyzed in 2 independent replicates. **H**. Western blot image of astrocytes pre-treated with MLi-2 before adding LLOME for 4 h. Membranes were probed for LC3 with ɑ-tubulin as a loading control. Histogram shows normalized LC3-II levels using two-way ANOVA with Tukey’s multiple comparison test comparing LLOME and MLi-2 treatments (FLLOME (1,8)= 67.28, *p*< 0.0001; FMLi-2 (1,8)= 0.4679, *p*= 0.5133; Finteraction(1,8)= 1.905, *p*= 0.2048). Data are means ± SEM from 3 independent replicates. **I**. Working model showing that LRRK2’s role in ruptured lysosomes is independent of lysophagy. Scale bar= 20 µm.

One widely reported effect of lysosome membrane disruption is the induction of lysophagy, a mechanism to clear ruptured lysosomes from cells via selective autophagy. Galectin-3 (Gal3) is mostly distributed in a diffuse fashion in the cell in normal conditions, but is recruited to the ruptured membrane and initiates lysophagy after a high degree of lysosomal membrane damage(*34*). In our experimental paradigm, Gal3 was indeed recruited to a subset of lysosomes after LLOME treatment (Fig. 2D, E). However, LRRK2-positive LYS were largely Gal3 negative (Fig. 2D, F), suggesting that LRRK2:JIP4 are recruited to a different lysosomal pool than those which will be degraded by lysophagy.

To further determine if LRRK2 modifies lysophagy, we pre-treated cells with the potent LRRK2 kinase inhibitor MLi-2 before adding LLOME. MLi-2 was able to completely block endogenous LRRK2 kinase activity in mouse primary astrocytes as observed by inhibition of both the LRRK2 auto-phosphorylation site pS1292 and of pT73 on RAB10, a known substrate of LRRK2 (Fig. S2F). In contrast, LRRK2 kinase inhibition did not affect Gal3 recruitment to the LYS in the presence of LLOME (Fig. 2G) and did not modify the autophagic response triggered by LLOME, as measured by LC3 lipidation (Fig. 2H). Taken together, our data suggest that LRRK2 plays a lysophagy-independent role in the membrane of ruptured lysosomes (Fig. 2I). We therefore considered whether LRRK2 recruitment occurs before the lysosomal membrane is ruptured enough to recruit Gal3 and trigger lysophagy and whether JIP4 might be important in mediating effects of LRRK2 on lysosomal function.

### LRRK2 recruits JIP4 to ruptured lysosomes in a kinase-dependent manner

As expected, JIP4 translocates to the membrane of LRRK2-positive LYS in a time-dependent manner (Fig. 3A). However, the response of JIP4 to LLOME was slower than that of LRRK2, with a maximal recruitment at 6 h instead of 2 h for LRRK2. It is also noteworthy that we noted the presence of several LRRK2-positive/JIP4-negative lysosomes, even after 6 h of LLOME exposure (Fig. 3A). MLi-2 did not reduce the lysosomal localization of LRRK2 (Fig. S3A), whereas completely blocked the recruitment of JIP4 to LRRK2-positive LYS (Fig. 3B), demonstrating that LRRK2 kinase activity is required for JIP4 translocation to the lysosomal membrane. JIP4 translocation was confirmed without transfection with LRRK2 (Fig. 3C, Fig. S3B), thus showing that LRRK2 is able to recruit JIP4 while expressed at endogenous levels. Endogenous LRRK2 pharmacological kinase inhibition was also able to arrest JIP4 lysosomal membrane localization (Fig. 3C). Notably, with endogenous LRRK2, the recruitment of JIP4 to the lysosomal membrane occurs 10 h after LLOME addition (Fig. S3B). These results demonstrate that increasing expression of LRRK2 accelerates JIP4 recruitment to membrane damaged lysosomes in a kinase-dependent manner.

**Figure 3.**
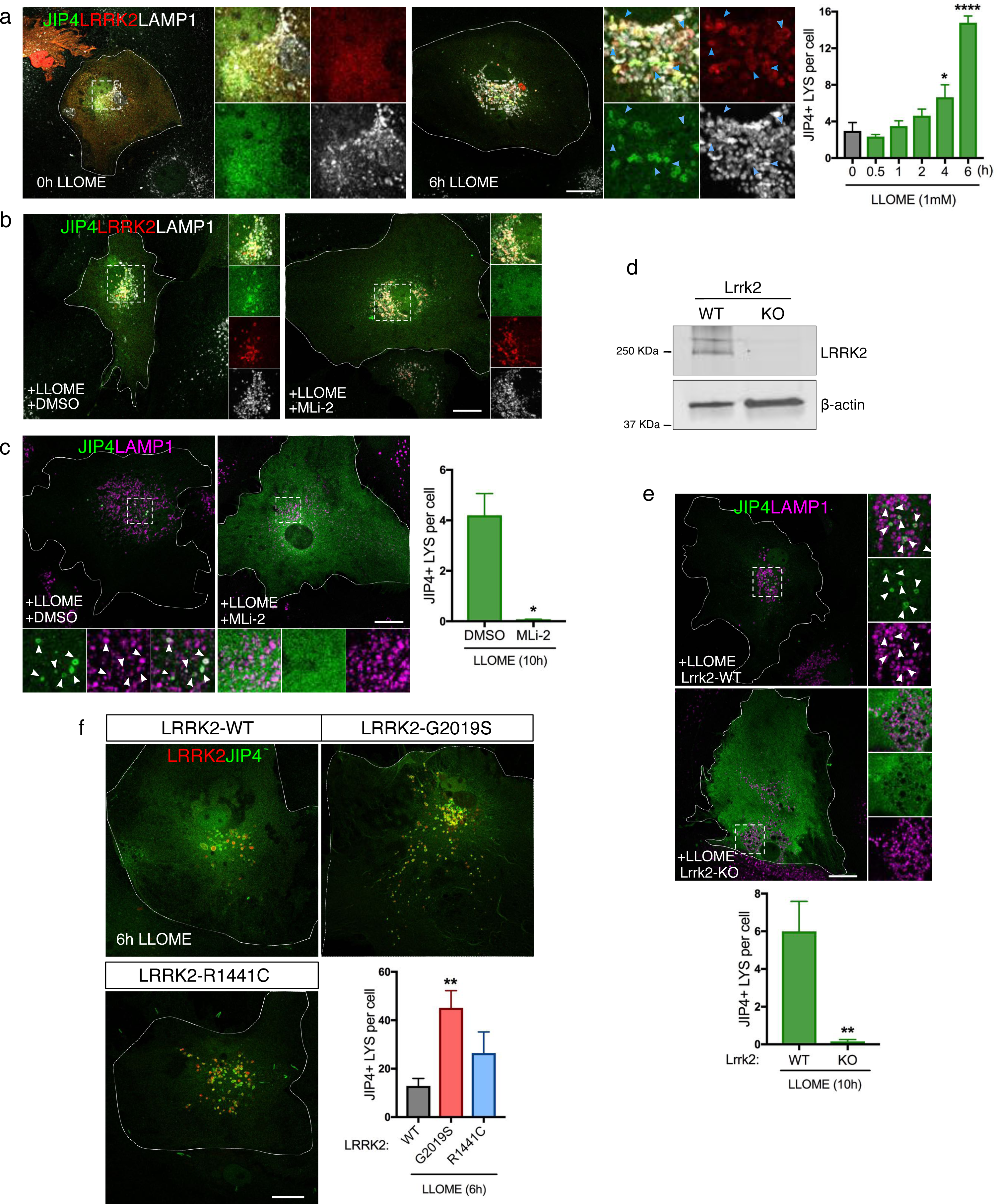
LRRK2 recruits JIP4 through its kinase activity. **A**. Representative confocal images of astrocytes expressing 3xflag-LRRK2 (red) and GFP-JIP4 (green), stained for LAMP1 (white) with or without treatment with LLOME. Histogram depicts the number of JIP4-positive lysosomes per cell after LLOME treatment from 3 independent replicates. Data are means ± SEM, *n*= 20 cells per condition. One-way ANOVA, F (5,11)= 41.4, *p*< 0.0001. Multiple comparisons were evaluated with Dunnett’s *post-hoc* test. **B**. Representative confocal images of astrocytes expressing 3xflag-LRRK2 (red) and GFP-JIP4 (green), stained for LAMP1 (white). Cells were pre-treated with DMSO or MLi-2 then incubated with LLOME. White arrowheads show colocalization and blue arrowheads indicate LRRK2-positive/JIP4-negative lysosomes. Scale bar= 20 µm. **C**. Astrocytes expressing GFP-JIP4 (green) and stained for LAMP1 (magenta) were pre-treated with DMSO or MLi-2 before adding LLOME. Histogram shows the number of JIP4-positive lysosomes per cell (*n*= 20-30 cells per condition). Data are means ± SEM from 3 independent replicates, unpaired *t-test* with Welch’s correction for unequal variance (*, *p*= 0.0407). **D**. Western blot confirming lack of LRRK2 expression in KO astrocytes compared to WT. **E**. Images of LrrK2-WT or Lrrk2-KO astrocytes transfected with GFP-JIP4 (green) and treated with LLOME. Statistical analysis used an unpaired *t-test* with Welch’s correction for unequal variances (**, *p*= 0.0032, *n*= 13 cells per condition). **F**. Astrocytes expressing GFP-JIP4 (green) were co-transfected with 3xflag-LRRK2-WT, 3xflag-LRRK2-G2019S or 3xflag-LRRK2-R1441C (red) and treated with LLOME. Histogram shows the number of JIP4-positive lysosomes per cell. Data are means ± SEM from 3 independent replicates, *n*= 20 cells per group. One-way ANOVA with Dunnett’s *post hoc* was applied, F (2,6)= 5.776, *p*= 0.0399.

To make sure these observations were not due to off-target effects of MLi-2, we compared the amount of lysosomal JIP4 in Lrrk2-WT and Lrrk2-KO astrocytes in the presence of LLOME. Consistent with prior data, cells lacking endogenous LRRK2 did not recruit JIP4 to ruptured lysosomes (Fig. 3D, E). Conversely, cells expressing the G2019S mutation (Fig. S3C) show a ∼3-fold increase in the number of JIP4-positive LYS per cell compared to cells only expressing the WT form of the protein (Fig. 3F).

### LRRK2 phosphorylates RAB10 and RAB35 in the lysosomal membrane

Since JIP4 recruitment to the lysosomal membrane is slower than LRRK2, we speculated that LRRK2 might require intermediate partners to recruit JIP4 upon lysosomal membrane damage. LLOME addition also triggers LRRK2-lysosomal localization in HEK293FT cells (Fig. S3D), so we performed an APEX2 screening in these cells in the presence or absence of LLOME (Fig. 4A). Among the potential candidates, we found a high number of endolysosomal markers, such as LAMP2, LAMTOR2, PSAP, RAB25 and GABARAPL1/2, confirming by proteomics the enrichment of LRRK2 in the endolysosomal system after LLOME treatment (Fig. 4B). Interestingly, two known substrates of LRRK2, RAB35 and RAB10, were also enriched by LLOME treatment (Fig. 4B). By staining for endogenous RAB35 in primary astrocytes, we observed a LLOME-driven recruitment of RAB35 to LRRK2-positive LYS (Fig. 4C), with similar results for a GFP tagged version of RAB10 (Fig. 4D). We therefore asked whether lysosomal membrane permeabilization triggers a LRRK2-dependent phosphorylation of both RAB proteins. LLOME addition induces a strong increase in phospho-RAB10 (pT73), that is not seen in cells expressing a kinase dead (K1906M) mutant LRRK2 or in cells treated with MLi-2 (Fig. 4E). Since there are no commercially available antibodies for phospho-RAB35, we used phos-tag gels that allow the recognition of phosphorylated forms of proteins by reducing their motility in the acrylamide gel. Cells treated with LLOME show a ∼3-fold increase in phospho-RAB35 expression (Fig. 4F), which was also blocked by co-incubation with MLi-2. Of interest, the LRRK2 auto-phosphorylation site pS1292 was not sensitive to the addition of LLOME. To make sure that LRRK2-mediated RAB phosphorylation occurs in the lysosomal membrane, we exogenously expressed LRRK2 and RAB10 in the presence of LLOME and stained also for phospho-RAB10 using the RAB10-pT73 antibody. As expected, the RAB10-pT73 signal colocalizes with LRRK2 in ruptured lysosomes (Fig. 4G). Collectively, these results show that lysosomal membrane damage triggers increased kinase activity towards RAB substrates.

**Figure 4.**
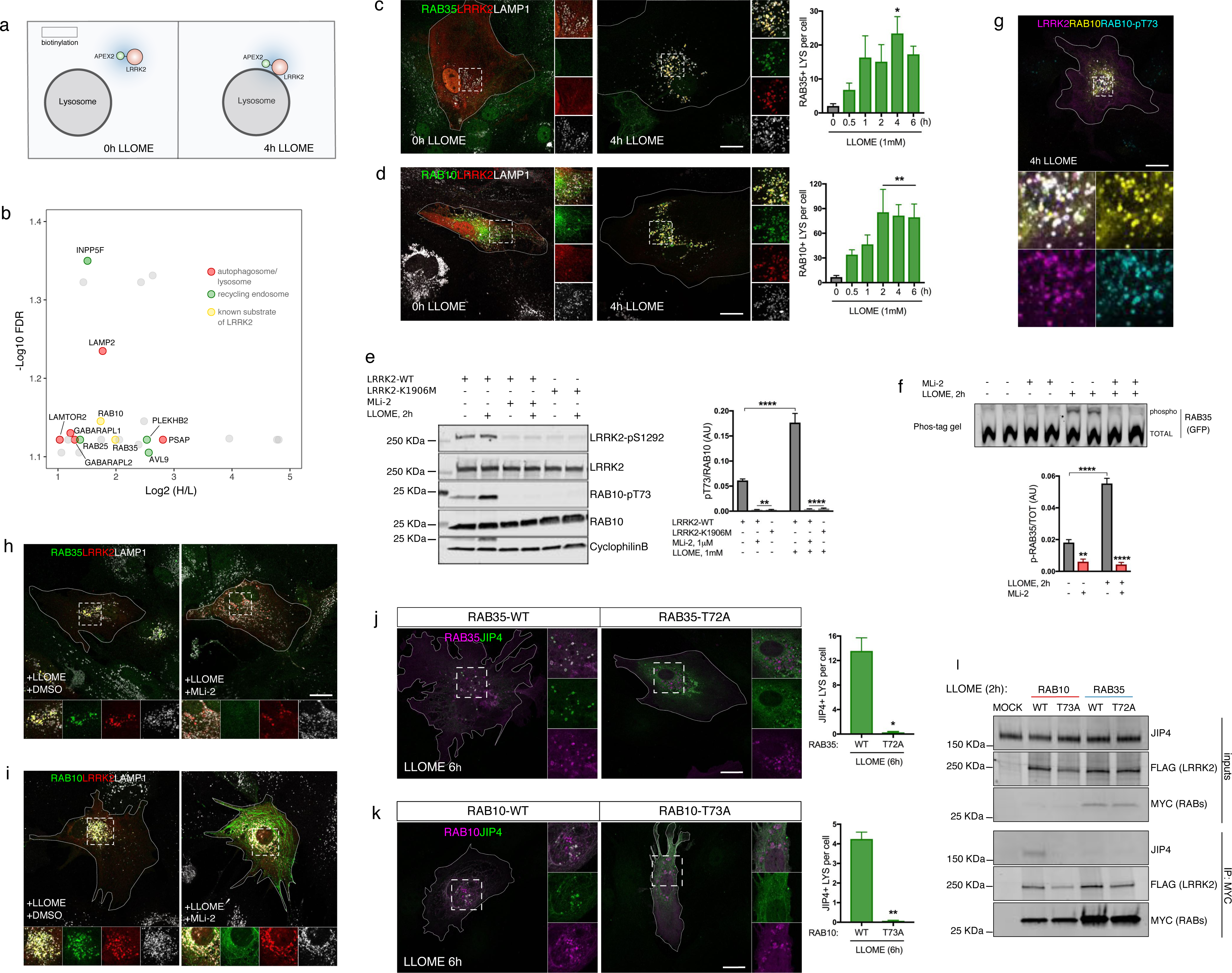
LRRK2 phosphorylates RAB35 and RAB10 in the membrane of ruptured lysosomes. **A**. Cartoon of the proximity ligation screening in HEK293FT cells in the presence or absence of LLOME. **B**. Volcano plot showing proteins in proximity with LRRK2 when cells were treated with LLOME showing the subset of proteins with a fold change > 2 and an FDR corrected *p*< 0.08 were considered, *n*= 3. **C**. Representative confocal images of astrocytes expressing 3xflag-LRRK2 (red), RAB35 (green) and LAMP1 (white) treated or note with LLOME. The number of RAB35-positive lysosomes per cell was quantified from 3 independent replicates with *n*= 20 cells per condition. Data are means ± SEM. One-way ANOVA with Dunnett’s multiple comparison test was applied, F (5,12)= 3.554, *p*= 0.0334. **D**. Confocal images of astrocytes expressing 3xflag-LRRK2 (red), GFP-RAB10 (green) and LAMP1 (white), either untreated or treated with LLOME. The number of RAB10-positive lysosomes per cell was evaluated by one-way ANOVA with Dunnett’s *post hoc* test, F (5, 114)= 4.392, *p*= 0.0011. Data are means ± SEM, *n*= 20 cells per condition. **E**. Representative western-blot images of HEK293FT cells transfected with 3xflag-LRRK2 (WT or K1906M) and pre-treated with DMSO or MLi-2 before LLOME was added. The normalized levels of RAB10-T73, LRRK2-pS1292 was measured. Data are means ± SEM in 4 independent replicates. Two-way ANOVA with Tukey’s multiple comparisons test was used to analyze the effect of treatment (LLOME) and kinase activity on RAB10-T73 expression, FLLOME (1,11)= 34.39, *p*= 0.0001; Fkinase (2,11)= 134.6, *p*< 0.0001; Finteraction (2,11)= 32.68, *p*< 0.0001. **F**. Phos-tag gel images of HEK293FT cells expressing 3xflag-LRRK2 and GFP-RAB35, pre-treated with DMSO or MLi-2 and incubated with LLOME. Normalized phospho-RAB35 levels were measured from 4 independent replicates. Data are means ± SEM. Two-way ANOVA with Tukey’s *post hoc*, FLLOME (1,12)= 69.38, *p*< 0.0001; FMLi-2 (1,12)= 218.9, *p*< 0.0001; Finteraction (1,12)= 83.62, *p*< 0.0001. **G**. Confocal picture of an astrocyte expressing 3xflag-LRRK2 (magenta), 2xmyc-RAB10 (yellow) and RAB10-T73 (cyan). **H-I**. Representative confocal images of astrocytes expressing 3xflag-LRRK2 (red), RAB35 (green, **H**), GFP-RAB10 (green, **I**) and LAMP1 (white) pre-treated with DMSO or MLi-2 and incubated with LLOME. **J**. Astrocytes were transfected with GFP-JIP4 and 2xmyc-RAB35-WT or 2xmyc-RAB35-T72A, and 48 h treated with LLOME for 6 h. Then, cells were fixed and stained for GFP (green) and myc (magenta). The number of JIP4-positive lysosomes per cell was quantified in both groups (*n*= 20 cells per condition), using an unpaired *t-test* with Welch’s correction (*, *p*= 0.0248). Data are means ± SEM from 3 independent replicates. **K**. Astrocytes were transfected with GFP-JIP4 and 2xmyc-RAB10-WT or 2xmyc-RAB10-T73A, and 48 h treated with LLOME for 6 h. Then, cells were fixed and stained for GFP (green) and myc (magenta). The number of JIP4-positive lysosomes per cell was quantified in both groups (*n*= 20 cells per condition), using an unpaired *t-test* with Welch’s correction (**, *p*= 0.0065). Data are means ± SEM from 3 independent replicates. **L**. HEK293FT cells were either mock transfected or transfected with 3xflag-LRRK2 along with 2xmyc-RAB10-WT, 2xmyc-RAB10-T73A, 2xmyc-RAB35-WT or 2xmyc-RAB35-WT. Cells were treated with LLOME (2 h), then lysates were subjected to immunoprecipitation (IP) with anti-myc antibodies and immunoblotted for endogenous JIP4, FLAG and MYC. White arrowheads indicate colocalization. Scale bar= 20 µm.

### LRRK2 recruitment of JIP4 occurs through RAB35 and RAB10 phosphorylation

Next, we questioned if LRRK2 could also be able to recruit RAB35 and RAB10 to the lysosomal membrane in a kinase-dependent manner. LRRK2 pharmacological kinase inhibition lead to a complete blockage of RAB35 and RAB10 recruitment to LRRK2-positive LYS (Fig. 4H, I). In astrocytes expressing LRRK2 at endogenous levels, both RAB35 and RAB10 were recruited to the lysosomal membrane after LLOME treatment (Fig. S4A, B), which was blocked by MLi-2 (Fig. S4C, D). These results show that endogenous LRRK2 is able to relocalize both RAB proteins to permeabilized LYS in a kinase-dependent manner. As both RAB35 and RAB10 are recruited to the LYS in a similar fashion to LRRK2, we wanted to determine if RAB35 or RAB10 were important to maintain LRRK2 at the lysosomal membrane. However, depletion of RAB35 and RAB10 expression (Fig. S4E, F) did not alter the ability of LRRK2 to translocate to ruptured lysosomes (Fig. S4G). Taken together, our data indicates that LRRK2 precedes and mediates the recruitment of RAB35 and RAB10 in a kinase-dependent fashion.

In astrocytes transiently transfected with tagged versions of LRRK2, JIP4 and RAB35, and stained for endogenous RAB10, all four proteins are present in the same structures (Fig. S4H), suggesting a possible link between the two RAB proteins and JIP4. JIP4 has been previously shown to require several RAB GTPases to ensure its presence in the recycling endosomal membrane(*35*), so we hypothesized that LRRK2 is able to recruit JIP4 to ruptured lysosomes through the phosphorylation of RAB35 and RAB10. First, we examined the response to LLOME of RAB35/RAB10 mutants that cannot be phosphorylated by LRRK2 (T72/73A) compared to their WT counterparts. Both RAB phospho-null mutants showed a significant decrease in their lysosomal localization after LLOME treatment, with a diffuse distribution in the cytosol. RAB35-T72A mutant displayed a ∼50% reduction compared to the WT form while RAB10-T73A showed an ∼80% decrease compared to RAB10-WT (Fig. S4I, J). Next, we wanted to know if the phospho-null version of RAB GTPases affects the recruitment of JIP4 to ruptured LYS. In both cases, cells expressing the phospho-null mutation were unable to recruit JIP4 to the lysosomal membrane, even in cells where RAB35-T72A and RAB10-T73A had a lysosomal localization (Fig. 4J, K). Using co-immunoprecipitation in cells treated with LLOME, we were able to see endogenous JIP4 physically interacting with RAB10-WT but not RAB10-T73A (Fig. 4L). Under the same conditions, we failed to detect a physical interaction between JIP4 and RAB35. Taken together, our data show that LRRK2 recruits JIP4 through the phosphorylated forms of RAB35 and RAB10, and JIP4 is a RAB effector in the context of lysosomal membrane damage.

### JIP4 enhances the formation of tubular structures emanating from lysosomes

To gain more insight into the role of the JIP4 in the lysosomal membrane, we imaged primary astrocytes transfected with LRRK2 and JIP4 using an Airyscan detector. The improved resolution of this microscopy approach allowed us to observe the presence of JIP4-positive/LRRK2-negative/LAMP1-negative tubular structures that stem from LYS (Fig. 5A-B, Suppl movie 1). Consistent with the observation that G2019S mutation leads to higher recruitment of JIP4 to the LYS, this mutation was associated with a higher number of tubules (Fig. 5C). JIP4-positive lysosomal tubules were also negative for LAMP1 when imaging living cells (Fig. S5A). Furthermore, these tubules were negative for LIMP2, another typical lysosomal membrane marker (Fig. S5B) and for the lysosomal luminal marker Dextran-555 (Fig. S5D). Next, we analyzed the tubular structures using FIB-SEM on astrocytes. First, we observed that the JIP4/LAMP1-positive compartments presented electron-dense structures (Fig. 5D, S5C), consistent with their lysosomal nature. By analyzing the JIP4-positive tubules at the structural level, we were able to confirm that the tubular membrane originates from the lysosomal membrane, albeit being negative for LAMP1 (red arrowheads in Fig. 5D and Suppl Fig. 5C-D). JIP4-positive lysosomal tubules display different morphologies, in that they dramatically differ in their length and thickness (Fig. 5D, S5C-D, suppl movie 6). Of note, we could see the ER surrounding the JIP4-positive tubule (Fig. 5E in magenta) and a microtubule connecting with the tubule (Fig. 5D, blue arrowheads and Fig. 5E in cyan), suggesting a role of these compartments in tubule dynamics.

**Figure 5.**
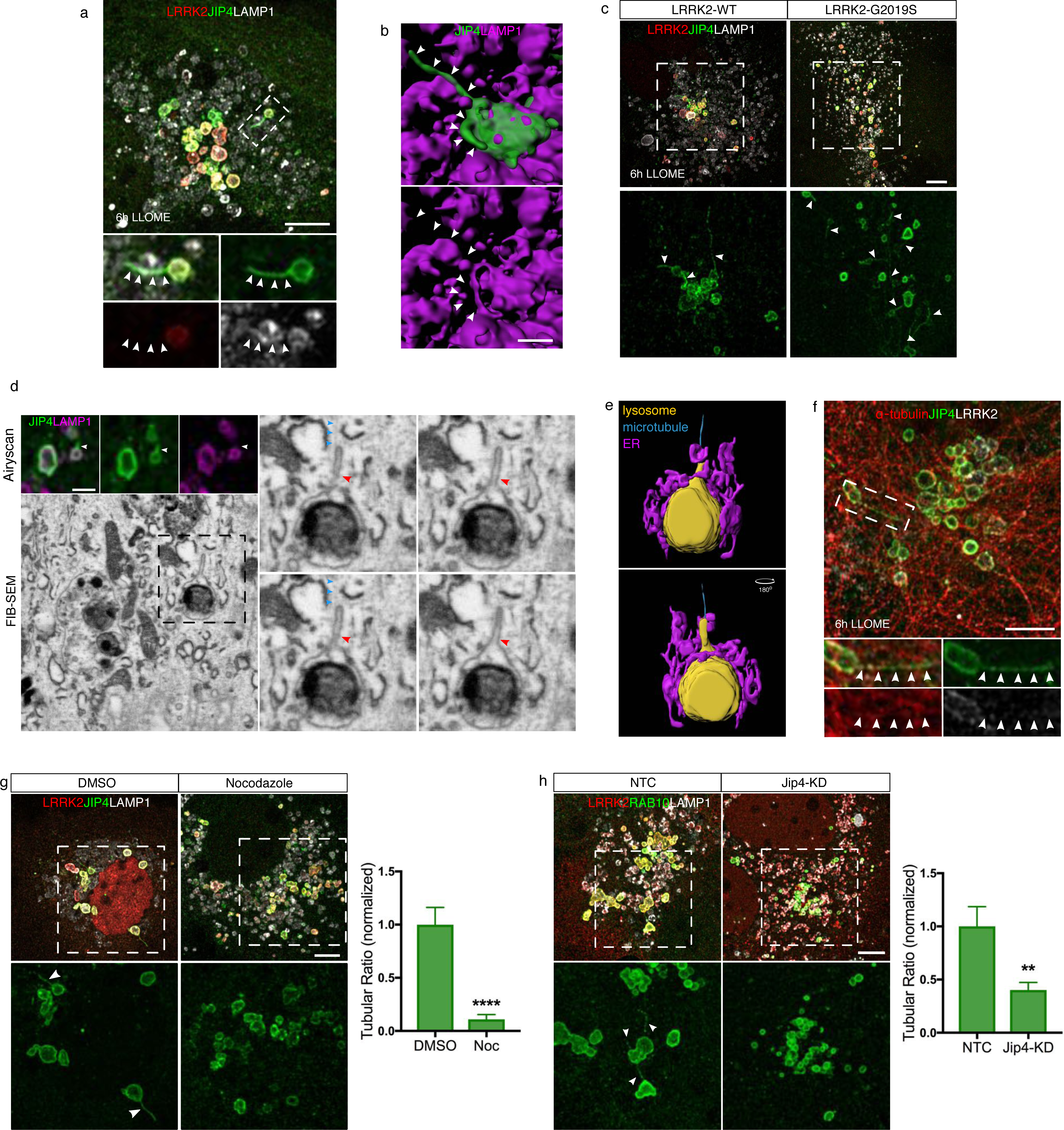
JIP4 promotes the formation of LAMP1-negative lysosomal tubular structures. **A**. Representative airyscan image of an astrocyte expressing GFP-JIP4 (green), 3xflag-LRRK2 (red) and LAMP1 (white), with LLOME. **B**. 3D surface reconstruction using Imaris was done in astrocytes transfected with 3xflag-LRRK2, mNeonGreen-JIP4 (green) and LAMP1-HaloTag (magenta). **C**. Confocal images processed with the airyscan module comparing the number of JIP4-positives tubules in cells expressing LRRK2-WT and LRRK2-G2019S. **D**. FIB-SEM image of a tubule stemming from a lysosome in a 3xflag-LRRK2-G2019S transfected astrocyte. Upper panel shows the airyscan image of two LAMP1-HaloTag (magenta)/mNeonGreen-JIP4 (green) labeled lysosomes. Lower panel shows the correlated electron microscopy image and right panel shows a cropped lysosome with a tubule in different z planes. **E**. 3D surface reconstruction of (D) showing a microtubule (cyan) and the ER (magenta) in contact with the lysosome (yellow). **F**. Super-resolution image of an astrocyte expressing 3xflag-LRRK2-G2019S (white), GFP-JIP4 (green) and ɑ-tubulin (red). **G**. Representative super-resolution images of astrocytes expressing 3xflag-LRRK2-G2019S (red), GFP-JIP4 (green) and LAMP1 (white) and treated with LLOME. 2 h before fixation, DMSO or 10 µM Nocodazole was added. JIP4 tubulation index was assessed between the two groups (DMSO and Nocodazole) using an unpaired *t-test* with Welch’s correction (****, *p*< 0.0001), *n*= 36-38 cells. Data are means ± SEM from 3 independent replicates. **H**. Confocal super-resolution images of astrocytes expressing 3xflag-LRRK2-G2019S (red), RAB10 (green) and LAMP1 (white) and incubated with LLOME for 10 h. RAB10 tubulation index was assessed between the two groups (NTC and JIP4-KD) using an unpaired *t-test* with Welch’s correction (**, *p*= 0.0041), *n*= 41 cells. Data are means ± SEM from 2 independent replicates. White and red (FIB-SEM picture) arrowheads indicate lysosomal tubular structures, and blue arrowheads indicate microtubules. Scale bar= 5 µm or 2 µm (**B** and **D**).

It has been shown that JIP4, through its interaction with motor proteins and microtubules, is required for the formation of endosomal sorting tubules(*36*). Consistent with the FIB-SEM observations, we saw JIP4-positive tubules that co-localized with ɑ-tubulin (Fig. 5F) and disruption of microtubules with nocodazole was association with a prevention of tubules formation (Fig. 5G), indicating that these structures are dependent on microtubules. Endogenous RAB10 but not RAB35 was present in a subset of JIP4-positive tubules (Fig. S5E, F) (consistent with the Co-IP results obtained in Fig. 4L), which allowed us to use RAB10 to determine if JIP4 was necessary for tubule formation. Overexpression of JIP4 lead to an increase in the number of tubules (Fig. S5G), and cells knocked-down for JIP4 (Fig. S5G) showed fewer tubules (Fig. 5H). Therefore, JIP4 is required for the extension of lysosomal membranes into tubules, likely via an interaction with microtubules.

### JIP4-positive tubules release vesicular structures that interact with other lysosomes

We next used super-resolution imaging to observe tubular dynamics in living cells, without fixation. JIP4-positive tubules bud, extend and release from lysosomes to form vesicular structures that were released into the cytosol (Fig. 6A, E, Suppl movie 2, Suppl movie 3). Of note, JIP4-positive vesicles can have several different behaviors, in that the scission can occur in the base of the tubule or from the tip (Fig. 6A, Suppl movie 2). Alternatively, tubules can retract into a vesicle that is ejected to the cytosol (Fig. 6A, E, Suppl movie 2, Suppl movie 3). Although JIP4-positive vesicles are motile in the cytosol, we often detected stationary vesicles in contact with other lysosomes (Fig. 6B). It was also possible to identify moving vesicles that stop to interact with a lysosome transiently, and then be released to move elsewhere (Suppl movie 4). Furthermore, using FIB-SEM, we were able to find a JIP4-positive vesicle contacting a lysosome (red arrowhead, Fig. 6C). The recipient lysosomes seem to be active, as we can observe these interactions happening in MR-CTSB-positive lysosomes in long periods of time (almost 4 min, white arrowhead in Fig. 6D and Suppl movie 5). Finally, we occasionally observed a resolved tubule forming a vesicle that is then able to interact with other lysosomes (Fig. 6E, Suppl movie 3). Taken together, our data identify that ruptured LYS form JIP4-positive tubules to release vesicular structures that can then contact other lysosomes.

**Figure 6.**
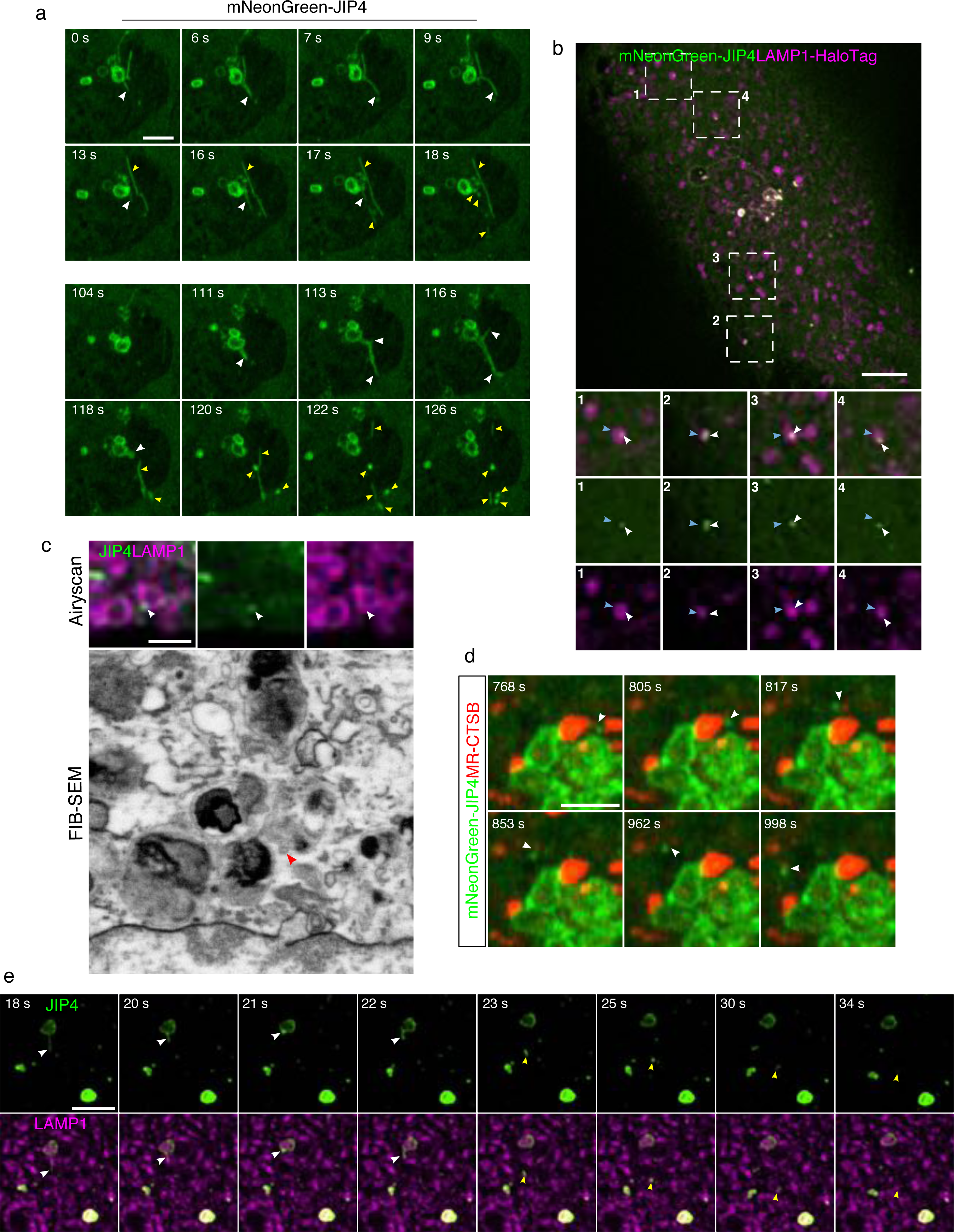
JIP4-positive tubules are resolved in vesicular structures that contact with other lysosomes. **A**. Time-lapse fast airyscan confocal images of an astrocyte expressing 3xflag-LRRK2-G2019S and mNeonGreen-JIP4 (green) showing a group of JIP4-positive lysosomes for over 2 min, where the different tubular dynamics were under display (15 slices/ frame were taken at 1 frame/ second). **B-C**. Astrocytes expressing 3xflag-LRRK2-G2019S, mNeonGreen-JIP4 (green) and LAMP1-HaloTag (magenta) were analyzed with a SorA microscope to perform spinning disk super-resolution microscopy. **B**. A single frame of an astrocyte where several JIP4-positive vesicles (white arrowhead) are in close proximity to other lysosomes (blue arrowhead). **C**. Time-lapse confocal images of a resolved tubule that after ejection to the cytosol contacts other lysosomes (15 slices/ frame, 1 frame/ second). **D**. FIB-SEM image of a JIP4-positive vesicle associated with a lysosome in a 3xflag-LRRK2-G2019S transfected astrocyte. Upper panel shows the airyscan image of a group of lysosomes (LAMP1, magenta) and a vesicle labeled with JIP4 (green, white arrow). Lower panel shows the correlated electron microscopy image with a red arrowhead pointing the vesicle. **E**. LLOME-treated astrocytes expressing 3xflag-LRRK2-G2019S, mNeonGreen-JIP4 (green) and stained with Magic-Red CTSB (red) were analyzed with a fast airyscan confocal microscope for almost 4 min, at 6.05 seconds/ frame. White arrowhead marks a JIP4-positive vesicle associated with an active lysosomes (red). White arrowheads indicate JIP4-positive lysosomal tubules, yellow arrowheads show resolved tubules (vesicular structures and scissioned tubules) (**A**-**C**). Scale bar: 2 µm (**D**), 2.5 µm (**A**, **C**, **E**), 5 µm (**B**).

In summary, LRRK2 gets recruited to permeabilized LYS that are not sufficiently damaged to trigger lysophagy. At the lysosomal membrane, LRRK2 is able to phosphorylate RAB35 and RAB10, leading to their retention in the membrane. This event is required to bring the motor adaptor protein JIP4 to the lysosomal membrane where it helps forming tubular structures along microtubules that can secondarily form small vesicles that can interact with other lysosomes (Fig. S5H). We call this newly described process LYsosomal Tubulation/sorting driven by LRRK2 or LYTL.

## DISCUSSION

Although a genetic link between LRRK2 and PD was first reported in 2004(*1*),(*37*), the role of this kinase in the cell remains uncertain. The localization of LRRK2 to intracellular membranes(*38*),(*39*),(*40*) and the observation that RAB GTPases are kinase substrates for LRRK2 (*8*),(*41*),(*9*), suggests that LRRK2 might be involved in vesicle-mediated transport(*42*). However, the major phenotype seen in LRRK2 knockout animals or in animals treated with LRRK2 kinase inhibitors is accumulation of lysosomal damage. This suggests to us that a major role for endogenous LRRK2 must be related to lysosomes.

Here we describe how LRRK2 can be recruited to the membrane of partially damaged and permeabilized lysosomes to trigger a newly described process that we call LYTL. A striking feature of LYTL is the activation of LRRK2 kinase activity. This is reminiscent of activation at other membranes, including at the *trans*-Golgi network (TGN) (*8*),(*43*).

However, that observation depends on overexpression of RAB29 whereas LYTL requires only modest lysosomal membrane damage, at levels less than those required to trigger lysosphagy. Consequentially to activation, LRRK2 phosphorylates RAB35 and RAB10, leading to their retention in the lysosomal membrane likely in turn due to diminished binding of p-RABs to GDI1/2(*21*). At the same time, p-RABs are able to recruit JIP4 to membrane damaged lysosome, consistent with prior data showing enhanced binding of p-RABs to this motor adaptor protein(*10*). Thus, JIP4 is a novel RAB effector in the lysosomal membrane, similar to the role previously described for JIP4 in recycling endosomes(*35*).

Sorting at other cellular compartments such as endosomes occurs through the formation of tubular structures that bud and extend from organelles. Tubular structures are then severed to produce vesicles that travel to the plasma membrane or the TGN(*44*). Our data show that JIP4 promotes the formation of such tubular structures at the lysosome(*36*). It is likely that JIP4 mediates tubule dynamics by recruiting motor proteins(*30*), since we have demonstrated that tubule extension requires microtubules. JIP4-positive tubules undergo scission to generate smaller vesicular structures that travel through the cytosol and interact with other lysosomes, although we cannot rule out the possibility of contact with other cellular organelles. Neither the tubules nor the resolved material are positive for lysosomal membrane markers (LAMP1 or LIMP2), and the JIP4-positive tubules were resistant to PFA fixation. These observations argue against LYTL being involved in proto-lysosome formation through lysosomal reformation (LR), since LR tubules are amenable to PFA fixation(*45*) and positive for LAMP1(*46*). To our knowledge, these observations are the first evidence of lysosomal sorting in mammalian cells.

We view the activation of LRRK2 by lysosomal membrane permeabilization as an analogous process to the activation of another PD-associated kinase, PINK1, by mitochondrial damage(*47*). In fact, one of the functions of the PINK1/Parkin system during mitochondrial stress is the release of mitochondria-derived vesicles to lysosomes(*48*),(*49*). There are still unanswered questions regarding LYTL, such as why ruptured lysosomes release membranous content and what the effect of vesicle contact with recipient lysosomes that will be the subject of future investigations.

We therefore identified a newly described cellular process that promotes lysosomal sorting material from permeabilized lysosomes. LYTL is controlled by LRRK2 kinase activity since the recruitment of all the downstream components is completely blocked by MLi-2. Conversely, the G2019S mutation in LRRK2 that is pathogenic for PD increases both the lysosomal localization of LRRK2 and JIP4 recruitment and tubulation. Considering the proposed centrality of lysosome biology in PD pathogenesis, it is possible that LYTL is involved in disease mechanisms.

## ACKNOWLEDGEMENTS

This research was supported in part by the Intramural Research Program of the NIH, National Institute on Aging (MRC) and the NIA Postdoctoral Funding Opportunity (LB-P). We thank Dr. David C. Gershlick (Cambridge University), Dr. Juan S. Bonifacino (NIH) and Dr. Dorien Roosen (FMP Berlin) for critical feedback, also Dr. Michael Schwake (Universitat Bielefeld) for kindly providing the LIMP2-myc plasmid. We also thank Eric Balzer (Nikon) for assistance using the SorA spinning-disk super-resolution microscope.

## AUTHOR CONTRIBUTIONS

Conceptualization, L.B-P., M.R.C.; Methodology, L.B-P., A.B., C.D.W., E.L., C.K.E.B., S.S.A., A.M, N.L., R.K.; Formal Analysis, L.B-P., M.R.C.; Investigation, L.B-P., A.B., E.L., Y.L., C.D.W., J.H.K.; Writing, L. B-P., M.R.C.; Supervision, M.R.C.; Funding, M.R.C., Y.L., L.B-P.

## DECLARATION OF INTERESTS

The authors have no competing interests related to this work. R.K. is currently an employee of Abcam.

## SUPPLEMENTARY MATERIAL

**SUPPLEMENTARY FIGURE S1. LRRK2 mutations and domain analysis affect the lysosomal recruitment of LRRK2. A.** Primary mouse cultures of cortical astrocytes, cortical neurons and microglia were made from *Lrrk2*-WT and *Lrrk2*-KO animals. RNA was extracted from them, as well as from microglia extracted from adult mouse brain. RNAseq was performed, and the mRNA expression of GFAP (astrocyte marker), MAPT (neuron marker), TMEM119 (microglia marker) and LRRK2 was plotted comparing Lrrk2-WT and KO cells. **B**. Schematic representation of LRRK2 and its domains, marking in orange the pathogenic hyperactive mutations (R1441C and G2019S) and in purple the artificial inactive mutations (T1348N and K1906M). **C**. Representative confocal images of astrocytes expressing 3xflag-LRRK2-WT, 3xflag-LRRK2-G2019S, 3xflag-LRRK2-K1906M and 3xflag-LRRK2-T1348N (red) and stained for LAMP1 (green). The number of LRRK2-positive lysosomes per cell was quantified and compared to the WT group in a one-way ANOVA with Dunnett’s multiple comparisons test of 3 independent replicates (*n*= 20 cells), F (4,10)= 57.39, *p*< 0.0001. Data are means ± SEM. **D**. Schematic representation of LRRK2 and its domains, marking the multi-domain plasmids used in this experiment in blue. **E**. Representative confocal images of astrocytes transfected with 3xflag-LRRK2-Full Length, 3xflag-LRRK2-ΔHEAT, 3xflag-LRRK2-ΔWD40, 3xflag-LRRK2-HEAT, 3xflag-LRRK2-ROC-COR-KIN and 3xflag-LRRK2-ROC-COR (red) and stained for LAMP1 (green). **F**. The number of LRRK2-positive lysosomes per cell was quantified comparing the Full-Length vector with the rest. Data are means ± SEM. One-way ANOVA with Dunnett’s multiple comparisons test of 3 independent replicates (*n*= 20 cells), F (6,14)= 32.64, *p*< 0.0001. **G**. Histogram showing the amount of LRRK2 translocation to the lysosomes comparing the domain constructs to each other. Data are means ± SEM. One-way ANOVA was used with Tukey’s *post-hoc* of 3 independent replicates (*n*= 20 cells), F (4,10)= 5.766, *p*= 0.0114. White arrowheads indicate colocalization between LRRK2 and LAMP1. Scale bar: 20 µm.

**SUPPLEMENTARY FIGURE S2. Additional information on the APEX2 screening for LRRK2-membrane partners. A**. Representative confocal images of HEK293FT cells expressing 3xflag-APEX2-LRRK2 (red) and 3xflag-APEX2-NES (red) and stained with Neutravidin (green) to detect biotinylation. **B**. Same experiment than **A**, but lysates were analyzed by western blot instead of confocal imaging. **C**. Full scatter plot of the two replicates of the APEX2-LRRK2 screening compared to APEX2-NES. In red are the proteins that passed the threshold of 2-fold increase in the LRRK2 group compared to NES in both replicates. **D**. BLOS1, MUTED, JIP3, STAM1 and BICD2 (green) were cloned in a GFP construct and expressed along 3xflag-LRRK2 (red). No colocalization was observed between these proteins and LRRK2 in the lysosomal membrane (yellow). **E.** HEK293FT cells were either mock transfected or transfected with 3xflag-tagged *E. coli* beta-glucuronidase (GUS) as negative controls or with 3xflag-LRRK2. Protein lysates were subjected to immunoprecipitation (IP) with anti-flag antibodies and immunoblotted for endogenous JIP4 or FLAG, showing that LRRK2 interacts with JIP4. **F.** Mouse primary astrocytes carrying the WT form or the G2019S mutation of LRRK2 were pre-treated with DMSO or MLi-2, lysed and immunoblotted for LRRK2-pS1292, LRRK2, RAB10-T73, RAB10 and cyclophilinB as a loading control. Scale bar: 20 µm.

**SUPPLEMENTARY FIGURE S3. Effect of MLi-2 and pathogenic hyperactive LRRK2 mutations in LRRK2 recruitment to lysosomes. A**. Representative confocal images of astrocytes expressing 3xflag-LRRK2 (red) and LAMP1 (white), pre-incubated with DMSO or MLi-2 and treated with LLOME. Histogram represents the number of LRRK2-positive lysosomes per cell. Data are means ± SEM. Unpaired *t-test* was applied in *n*= 19-20 cells (*p*= 0.2847). **B**. Representative confocal images of astrocytes expressing GFP-JIP4 (green) and LAMP1 (magenta), treated with LLOME. Histogram represents the number of JIP4-positive lysosomes per cell. Data are means ± SEM. One-way ANOVA with Dunnett’s *post hoc* for multiple comparisons was applied, F (4,93)= 3.398, *p*= 0.0122. **C**. Representative confocal images of astrocytes expressing 3xflag-LRRK2-WT, 3xflag-LRRK2-G2019S and 3xflag-LRRK2-R1441C (red) and stained for LAMP1 (white). The number of LRRK2-positive lysosomes per cell was measured in 20 cells per group (from 3 independent replicates). Data are means ± SEM. One-way ANOVA with Dunnett’s *post hoc* was used, F (2,57)= 4.769, *p*= 0.0122. **D**. Representative confocal images of HEK293FT cells expressing 3xflag-LRRK2 (red) and LAMP2 (green) and treated with LLOME for the indicated time points. Histogram refers to the number of LRRK2-positive lysosomes per cell. Data are means ± SEM. One-way ANOVA with Dunnett’s test for multiple comparisons was applied, F (5,284)= 14.42, *p*< 0.0001. *n*= 40-56 cells per condition. Scale bar: 20 µm.

**SUPPLEMENTARY FIGURE S4. Additional information on the link between LRRK2 and both RAB GTPases (RAB35 and RAB10) upon lysosomal membrane permeabilization. B**. Representative confocal images of astrocytes expressing RAB35 (green) and LAMP1 (magenta), treated or not with LLOME. Histogram depicts the number of RAB35-positive lysosomes per cell. Data are means ± SEM. Unpaired *t-test* with Welch’s correction was applied (***, *p*= 0.0005). **C**. Representative confocal images of astrocytes expressing EGFP-RAB10 (green) and LAMP1 (magenta), treated or not with LLOME. Histogram depicts the number of RAB10-positive lysosomes per cell. Data are means ± SEM. Unpaired *t-test* with Welch’s correction was used (**, *p*= 0.0033). **D**. Representative confocal images of astrocytes expressing RAB35 (green) and LAMP1 (magenta), pre-incubated with DMSO or MLi-2 in LLOME-treated cells. Histogram corresponds to the number of RAB35-positive lysosomes per cell. Data are means ± SEM. Unpaired *t-test* with Welch’s correction was applied (**, *p*= 0.0087), *n*= 30-61 cells, in 3 independent replicates. **E**. Representative confocal images of astrocytes expressing EGFP-RAB10 (green) and LAMP1 (magenta), pre-incubated with DMSO or MLi-2 in LLOME-treated cells. Histogram corresponds to the number of RAB10-positive lysosomes per cell. Data are means ± SEM. Unpaired *t-test* with Welch’s correction was applied (*, *p*= 0.0336), *n*= 19-20 cells, in 3 independent replicates. **F-G**. Astrocytes were exposed to NTC siRNA and Rab35 siRNA or Rab10 siRNA were lysed and immunoblotted for RAB35 (**F**) and RAB10 (**G**) and ɑ-tubulin as a loading control. **H**. Representative confocal images of astrocytes expressing 3xflag-LRRK2 (red) and LAMP1 (green) in cells knocked-downed for Rab10 and Rab35, using an NTC as negative control. Histogram shows the number of LRRK2-positive lysosomes per cell. Data are means ± SEM. One-way ANOVA with Dunnett’s *post-hoc* was used, F (2,6)= 0.1706, *p*= 0.8471. *n*= 20 cells in 3 independent replicates. **I**. Confocal image shows an astrocyte expressing 2xmyc-RAB35 (blue), GFP-JIP4 (green), RAB10 (red) and 3xflag-LRRK2 (white) in LLOME-treated cells. **J**. 3xflag-LRRK2 was transfected along 2xmyc-RAB35-WT or 2xmyc-RAB35-T72A and treated with LLOME for 4 h. FLAG (magenta) and myc (cyan) were imaged under the confocal microscope and the number of RAB35 lysosomal structures per cell was quantified (*n*= 12 cells per group). Data are means ± SEM. Unpaired *t-test* was used (*, *p*= 0.0108) in 3 independent replicates. **K**. 3xflag-LRRK2 was transfected along 2xmyc-RAB10-WT or 2xmyc-RAB10-T73A and treated with LLOME for 4 h. FLAG (magenta) and myc (cyan) were imaged under the confocal microscope and the number of RAB10 lysosomal structures per cell was quantified (*n*= 12 cells per group). Data are means ± SEM. Unpaired *t-test* was used (**, *p*= 0.002) in 3 independent replicates. White arrowheads show colocalization. Scale bar: 20 µm.

**SUPPLEMENTARY FIGURE S5. Additional characterization of the JIP4-positive tubular structures. A**. Airyscan live cell super-resolution image of an astrocyte expressing 3xflag-LRRK2-G2019S, mNeonGreen-JIP4 (green) and LAMP1-RFP (magenta), treated with LLOME. **B**. Representative super-resolution image of an astrocyte expressing 3xflag-LRRK2-G2019S (red), GFP-JIP4 (green) and LIMP2-myc (white) treated with LLOME. **C**. Representative FIB-SEM image of a thin JIP4-positive tubule in a 3xflag-LRRK2-G2019S transfected astrocyte. Upper panel shows the airyscan image of a JIP4-positive lysosome forming a thin tubule (white arrowhead). Lower panel shows the correlated electron microscopy image with a red arrowhead pointing the lysosomal tubule. **D**. An example of the morphological diversity of the JIP4-positive tubules revealed by FIB-SEM. Left picture shows the airyscan image of a JIP4-positive lysosomes forming five different tubules. Left picture shows the segmented 3D reconstruction from the correlated electron microscopy image with white arrowheads marking the different tubules. **E**. Astrocytes transfected with 3xflag-LRRK2-G2019S (white) and GFP-JIP4 (green). 24 h later, fixable Dextran-555 (red) was incubated for 6 h. After 3 washes, Dextran was chased for at least 18 h and cells were treated with LLOME. **F**-**G**. Astrocytes exogenously expressing 3xflag-LRRK2-G2019S (white) and GFP-JIP4 (green) were treated with LLOME for 6 h and stained for endogenous RAB10 (red) (**F**) and endogenous RAB35 (red) (**G**). **H**. Astrocytes were treated with NTC or Jip4 siRNA. Western Blot shows JIP4 expression. **I**. Confocal super-resolution images of astrocytes expressing 3xflag-LRRK2-G2019S (white), RAB10 (red) and MOCK or GFP-JIP4 (green) transfected. RAB10 tubulation index was assessed between the two groups (MOCK and JIP4-OE) using an unpaired *t-test* with Welch’s correction (**, *p*= 0.0059), *n*= 30 cells. Data are means ± SEM from 2 independent replicates. **J**. Schematic representation of our working model. White arrowheads indicate lysosomal tubular structures. Scale bar= 2 µm (**C**), 5 µm.

**SUPPLEMENTARY MOVIE 1. JIP4-positive tubules are negative for LRRK2 and LAMP1**. Astrocytes expressing 3xflag-LRRK2, GFP-JIP and stained for LAMP1 were treated with LLOME for 6 h. Super-resolution stack was taking using airyscan and 3D reconstruction was made using Imaris software. Video was acquired at 15 frames/ second.

**SUPPLEMENTARY MOVIE 2. Super-resolution live cell imaging on JIP4-positive tubules**. Confocal microscopy of JIP4-positive tubules budding, extending and releasing vesicular structures in a living astrocyte expressing 3xflag-LRRK2-G2019S and mNeonGreen-JIP4 (green) treated with LLOME for 6 h. Video was acquired at 1 second/ frame for 20 and 18 seconds, and played at a speed of 3 frames/ second). Video corresponds to Fig. 7A. White arrowheads indicate JIP4-positive lysosomal tubules, yellow arrowheads show resolved tubules (vesicular structures and scissioned tubules).

**SUPPLEMENTARY MOVIE 3. Released vesicular structure interacting with other lysosomes**. Confocal microscopy of JIP4-release vesicular structure contacting with lysosomes, in a living astrocyte expressing 3xflag-LRRK2-G2019S, LAMP1-HaloTag (magenta) and mNeonGreen-JIP4 (green) treated with LLOME for 6 h. Video was acquired at 1 second/ frame for 34 seconds and played at a speed of 3 frames/ second). Video corresponds to Fig. 7C. White arrowheads indicate JIP4-positive lysosomal tubules, yellow arrowheads show resolved tubule (vesicular structure).

**SUPPLEMENTARY MOVIE 4. Moving vesicle temporary interacting with a lysosome.** Confocal microscopy from the same cell as Fig. 7C and Suppl movie 2. 3xflag-LRRK2-G2019S, mNeonGreen-JIP4 (green) and LAMP1-HaloTag (magenta) were transfected in primary astrocytes and treated with LLOME for 6 h. Video was acquired at 1 second/ frame for 56 seconds and played at a speed of 6 frames/ second. White arrowheads show moving JIP4-positive vesicle.

**SUPPLEMENTARY MOVIE 5. Super-resolution live cell imaging on vesicle contacting an active lysosome**. Confocal microscopy of JIP4-release vesicular structure contacting an active lysosome, in a living astrocyte expressing 3xflag-LRRK2-G2019S, and mNeonGreen-JIP4 (green) treated with LLOME for 6 h and incubated with Magic-Red CTSB (red) for 30 min. Each frame was acquired after 6.05 seconds, for 230 seconds (and played at a speed of 8 frames/ second). White arrowhead indicates a JIP4-positive vesicular structure interacting with a MR-CTSB-positive lysosome.

**SUPPLEMENTARY MOVIE 6**. **FIB-SEM reveals the morphological heterogeneity of the JIP4-positive tubules**. FIB-SEM segmentation of a JIP4-positive lysosome (from Fig. S5D) forming five tubules with different size and thickness. Segmentation was performed with Amira software.

## METHODS

### Cell culture

All procedures with animals followed the guidelines approved by the Institutional Animal Care and Use Committee of the National Institute on Aging. Astrocyte cultures were prepared from C57BL/6J newborn (P0) pups. First, dissected mouse cortices were incubated in 1 ml/cortex Basal Medium Eagle (BME) (Sigma-Aldrich), containing 5 U of papain/ (Worthington) for 30 min at 37°C. Five μg of DNAse I was added to each cortex preparation, and brain tissue was dissociated into single cells. Cells were washed twice with 10 volumes of BME and counted. Astrocyte cultures were plated in DMEM media (Thermo Scientific), supplemented with 10% FBS (Lonza) into 75 cm^2^ tissue culture flasks. For the preparation of purified astrocyte cultures, 7-10 day primary cultures were vigorously shaken to detach microglia and oligodendrocytes. Culture purity was assessed with GFAP for astrocytes, and OLIG2 and IBA1 to exclude oligodendrocytes and microglia. Cultures had 70-90% of astrocytes in all experiments. Astrocytes were used from passage 2 to passage 3. HEK293FT cells were maintained in DMEM containing 4.5 g/l glucose, 2 mM l-glutamine, and 10% FBS at 37°C in 5% CO2. HEK293FT cells and primary neuronal cultures were seeded on 12 mm coverslips pre-coated with matrigel.

### Reagents

LLOME (Sigma-Aldrich, L7393) was diluted in DMSO and added at 1 mM at the indicated time points. Nocodazole (Sigma-Aldrich, M1404) was diluted in DMSO and added at 10 µM 2 h before fixation. Fixable Dextran-Alexa Fluor 555, 10 KDa (ThermoFisher, D34679) was incubated for 6 h (2.5 mg/ml). Cells were then washed 3 times with PBS, and fresh media was added to chase Dextran overnight (18-24 h) before treating astrocytes with LLOME. MLi-2 (Tocris, 5756) was used at 1 µM, 90 min before LLOME addition. Magic-Red CTSB was obtained from ImmunoChemistry Technologies (938) and incubated at 1/250 in cells for 30 min. Then cells were washed 3x with PBS and analyzed under the microscope. LysoTracker Red DND-99 (ThermoFisher, L7528) was added at 1 µM for 30 min before washing the cells 3 times with PBS and analyzed them using confocal microscopy. LAMP1-HaloTag transfected cells were incubated with the JF646 peptide (Promega, GA1121) at 200 nM for 15 min, cells were then washed 3x and incubated with fresh media before treated with LLOME.

### Antibodies

The following primary antibodies were used: mouse anti-FLAG M2 (Sigma-Aldrich, F3165, 1/500 for ICC and 1/10.000 for WB), rabbit anti-JIP4 (Cell Signaling, 5519, 1/1.000 for WB), rat anti-LAMP1 (DSHB, 1D4B, 1/100 for ICC), rat anti-LAMP2 (DSHB, 1/100 for ICC), mouse anti-LAMP2 (DSHB, H4B4, 1/100 for ICC), rat anti-FLAG (Biolegend, 637302, 1/100 for ICC), rabbit anti-RAB7A (Abcam, ab137029, 1/200 for ICC), rat anti-myc (Chromotek, 9e1-100, 1/500 for ICC), chicken anti-GFP (AvesLab, GFP-1020, 1/500-1/1.000 for ICC), mouse anti-GFP (Roche, 11814460001, 1/10.000 for WB), goat anti-CTSB (R&D Systems, AF965, 1/500 for ICC), mouse anti-ɑ-tubulin (Cell Signaling, 3873, 1/350 for ICC and 1/15.000 for WB), rabbit anti-RAB35 (Proteintech, 11329-2-AP, 1/100 for ICC and 1/1.000 for WB), rabbit anti-RAB10 (Abcam, ab237703, 1/100 for ICC and 1/1.000 for WB), rabbit anti-RAB10 (phospho-T73) (Abcam, ab23026, 1/100 for ICC and 1/1.000 for WB), rabbit anti-LRRK2 (Abcam, ab133474, 1/1.000 for WB), rabbit anti-LRRK2 (phospho S1292) (Abcam, ab203181, 1/1.000 for WB), rabbit anti-EEA1 (Cell Signaling, 3288, 1/100 for ICC), mouse anti-β-actin (Sigma-Aldrich, A5441, 1/15.000 for WB), rabbit anti-LC3B (Cell Signaling, 2775, 1/1.000 for WB), rabbit anti-cyclophilin B (Abcam, ab16045, 1/15.000 for WB).

For ICC, unless otherwise stated, the secondary antibodies were purchased from ThermoFisher. The following secondary antibodies were used: donkey anti-mouse Alexa-Fluor 568 (A10037, 1/500), donkey anti-rabbit Alexa-Fluor 488 (A-21206, 1/500), donkey anti-mouse Alexa-Fluor 568 (A-21202, 1/500), donkey anti-rat Alexa-Fluor 488 (A-21208, 1/500), donkey anti-goat Alexa-Fluor 488 (A-11055, 1/500), donkey anti-rabbit Alexa-Fluor 568 (A10042, 1/500), donkey anti-mouse Alexa-Fluor 647 (A-31571, 1/500), goat anti-rat Alexa-Fluor 647 (A-21247, 1/250-1/500). Donkey anti-chicken Alexa-Fluor 488 (703-545-155, 1/500) and donkey anti-rat Alexa-Fluor 405 (712-475-153, 1/100) were obtained from Jackson ImmunoResearch.

### Cloning

Constructs for 3x-flag-tagged LRRK2 full length and domains, and GUS have been described previously(*50*),(*8*). JIP3, JIP4 and BICD2 Gateway PLUS shuttle clones were obtained from GeneCopoeia. BLOS1, MUTED and STAM1 cDNA were obtained from Dharmacon. RAB35, RAB10 and APEX-NES cDNA were acquired from Addgene (Addgene#49552, Addgene#49472 and Addgene#49386)(*51*),(*52*),(*53*). cDNAs were amplified with PCR and cloned into pCR8/GW/TOPO vector (ThermoFisher). All of them were then transferred into pDEST vectors using Gateway technology (ThermoFisher).

Full-length LRRK2 was transferred into the pDEST-Emerald GFP and p3xflag-APEX2-DEST vectors. JIP3, STAM1, BICD2, BLOS1 and MUTED were transferred into the pDEST-53 plasmid. JIP4 was transferred into the pDEST-53 and the pDEST-NeonGreen vectors. RAB35 was transferred into the pDEST-53 and the pCMV-2xmyc-DEST vectors. RAB10 was transferred into the pCMV-2xmyc-DEST plasmid. 2xmyc-RAB10-T73A and 2xmyc-RAB35-T72A were made by directed mutagenesis from the WT vectors using the QuickChange Lightning kit (Agilent). EGFP-RAB10 (Addgene#49472)(*52*), EGFP-Gal3 (Addgene#73080)(*34*) and LAMP1-RFP (Addgene#1817)(*54*) were purchased from Addgene. The LAMP1-HaloTag construct was provided by the Bonifacino lab and the LIMP2-myc vector was a gift from Michael Schwake(*55*). All the primers used for cloning are in Suppl Table 3. All the expression constructs used in this study are summarized in Suppl Table 4.

### Transfection

Transient transfections of HEK293FT cells and astrocytes were performed using Lipofectamine 2000 and Stem reagents (ThermoFisher). HEK293FT cells were transfected for 24 h and mouse primary astrocytes were transfected for 48 h. For siRNAs, cells were transfected with the SMARTpool ON-TARGET (Dharmacon) plus scramble or Rab35, Rab10 or Jip4 siRNAs using Lipofectamine RNAiMAX (ThermoFisher) transfection reagent for astrocytes. Astrocytes were incubated with siRNA for a total of 4 days before fixation or lysis.

### APEX2 reaction

HEK293FT cells were plated in 150 cm^2^ flasks previously coated with matrigel and transfected with the appropriate vectors. 24 h later, 500 μM biotin-phenol was pre-incubated with cells for 30 min at 37°C. 1 mM H2O2 was added for 1 min while gently mixing and the cells were subsequently washed two times with quenching buffer (TBS supplemented with 1 mM CaCl2, 10 mM sodium ascorbate, 1 mM sodium azide, and 1 mM Trolox), one time with PBS and an extra two times with quenching buffer with one min per wash. Cells were collected in 10 ml of quenching buffer and centrifuged at 3000xg for 10 min. Cells were lysed in RIPA buffer (Pierce) supplemented with 10 mM sodium ascorbate, 1 mM sodium azide, 1 mM Trolox, 1 mM DTT, and protease inhibitors (Roche Complete). Samples were briefly sonicated, spun down at 10.000xg for 10 min, equal amounts of proteins were applied to streptavidin magnetic beads (Thermo), and incubated 1 h at RT on a rotator. Cells were then washed sequentially with KCl (1 M), Na2CO3 (0.1 M) and Urea (2 M) in 0.1 M TRIS buffer, with a final two washes with RIPA buffer. Beads were eluted by boiling in 45 µl of 2x protein loading buffer supplemented with 2 mM biotin and β-Mercaptoethanol for 10 min. Protein lysates were loaded on an acrylamide gel and run for 1 h at 150 V. The gel was then stained with GelCode Blue stain (ThermoFisher) for 30 min and washed with dH2O overnight.

### Mass Spectrometry analysis

Protein was quantified and the lysates were mixed 1:1, having a final number of 2-3 replicates. Lysates were separated on a 4–20% polyacrylamide gel (Bio-Rad). 8 bands were cut from each replicate. Gel bands were reduced with tris(2-carboxyethyl)phosphine (TCEP), alkylated with N-Ethylmaleimide (NEM), and digested with trypsin. The system used for data acquisition is an Orbitrap Lumos mass spectrometer (Thermo Fisher Scientific) coupled with a 3000 Ultimate high-pressure liquid chromatography instrument (Thermo Fisher Scientific). Extracted peptides were desalted then separated on an ES803 column (Thermo Fisher Scientific) using a gradient with mobile phase B (MP B) increasing from 5 to 27% over 60 min. The composition of MP A and MP B are 0.1% formic acid in 100% HPLC acetonitrile, respectively. The LC-MS/MS data were acquired in data-dependent mode. The survey scan was performed in orbitrap with the following parameters: mass range: 350-1500 m/z; resolution: 120k at m/z 400, AGC target 4 × 10e5 ions. The product scan was performed in ion trap with the following parameters: Collision-induced dissociation on as many precursor ions as allowed in 3 Sec; isolation window: 1.6 Da. Database search and ratio calculation were performed using Proteome Discoverer 2.2 software.

Conditions used in the database are listed below. Database: Sprot Human database; Modifications: 13C(6) (R), 13C(6) (K), Oxidation (M), NEM(C). H/L ratios are calculated for each sample with (Fig. 4) or without (Fig. 1) normalization.

### Phos-tag gels

HEK293FT cells were transfected with 3xflag-LRRK2 and GFP-RAB35 plasmids, and 24 h later treated with LLOME or DMSO for 2 h. Cells were then lysed in 10x Cell Lysis Buffer (Cell Signaling) supplemented with EDTA-free protease inhibitor (Roche) and 1x Halt phosphatase inhibitor. Lysates were cleared by centrifugation at 20.000xg. Phos-tag SDS-Page was performed following the vendors instructions (Wako). Briefly, gels for Phos-tag SDS/PAGE consisted of a stacking gel (4.5% acrylamide, 350 mM Bis-Tris/HCl, TEMED and 0.05% APS) and a separating gel (12% acrylamide, 350 mM Bis-Tris/HCl, 50 μM Phos-tag™ solution, 100 μM ZnCl2, TEMED and 0.05% APS). Lysates were electrophoresed at 80 V for the stacking gel and at 150 V for the separating. After SDS-PAGE, the gels were washed two times (20 min each) with transfer buffer containing 10 mM EDTA. Gels were transferred to membranes by semi-dry trans-Blot Turbo transfer system (Biorad).

### Confocal microscopy

Confocal images were taken using a Zeiss LSM 880 microscope equipped with a 63x1.4 NA objective. Super-resolution imaging was performed using the Airyscan mode. Raw data were processed using Airycan processing in ‘auto strength’ mode with Zen Black software version 2.3. Live cell super-resolution imaging was performed in Fast AiryScan mode on an inverted Zeiss LSM 880 Fast AiryScan microscope equipped with a 63x 1.4 NA objective, and environmental chamber to maintain cells at 37°C with humidified 5% CO2 gas during imaging. Immersion oil for 37°C conditions was used during imaging.

For spinning disk super-resolution microscopy we used a W1-SoRa super-resolution spinning disk microscope (Nikon) with a 60x 1.49 NA oil immersion objective and a 2.8x intermediate magnification (168x combined), with the offset micro-lensed SoRa disk, and environmental chamber to maintain cells at 37°C with humidified 5% CO2 gas during imaging. The deconvolution settings were: 10 iterations of Landweber decon.

The two colors were acquired simultaneously using a Cairn twin-cam emission splitter and two Photometrics prime 95b sCMOS cameras, a 565LP DM, and appropriate emission cleanup filters. Only cells with a healthy appearance and not showing obvious artifacts due to overexpression were imaged.

### Tubular ratio measurements

Astrocytes were transfected with the 3xflag-LRRK2-G2019S mutation plasmid, and co-transfected or not with the GFP-JIP4 vector. After fixation and staining, cells were imaged with a confocal microscope using the Airyscan module to enhance resolution. The tubular ratio in each cell was measured as: Ratio= #JIP4+tubules/ #JIP4+lysosomes. The same goes for endogenous RAB10. For each independent replicate, the ratio was normalized by the average value of the control group (i.e. DMSO, NTC and MOCK). Data were generated from cells pooled across at least two independent experiments. Only cells with 10 or more JIP4 or RAB10-positive lysosomes per cell were imaged.

### Co-Immunoprecipitation

HEK293FT cells transfected with 3xflag-LRRK2, 2xmyc-RAB10, 2xmycRAB10-T73A, 2xmyc-RAB35 and 2xmyc-RAB35-T72A plasmids were lysed in IP buffer (20 mM Tris-HCl pH 7.5, 300 mM NaCl, 1 mM EDTA, 0.3% Triton X-100, 10% Glycerol, 1x Halt phosphatase inhibitor cocktail (ThermoFisher) and protease inhibitor cocktail (Roche)) for 30 min on ice. Lysates were centrifuged at 4°C for 10 min at 20.000xg and supernatant further cleared by incubation with Easy view Protein G agarose beads (Sigma-Aldrich) for 30 min at 4°C (only for FLAG IP). After agarose beads removal by centrifugation, lysates were incubated with anti-flag M2 agarose beads (Sigma-Aldrich) or myc-Trap agarose beads (Chromotek) for 1 h at 4°C on a rotator. Beads were washed six times with IP wash buffer (20 mM Tris-HCl pH 7.5, 300 mM NaCl, 1 mM EDTA, 0.1% Triton X-100, 10% Glycerol) and eluted in 1X kinase buffer (Cell Signaling), containing 150 mM NaCl, 0.02% Triton and 150 ng/µl of 3xflag peptide (Sigma-Aldrich) by shaking for 30 min at 4°C (for FLAG IP) or by boiling samples in 2X loading buffer with β-Mercaptoethanol for 5 min (for MYC IP).

### Immunostaining

Primary cultures of astrocytes or HEK293FT cells were fixed with 4% PFA for 10 min, permeabilized with PBS/ 0.1% Triton for 10 min and blocked with 5% Donkey Serum for 1 h at RT. Primary antibodies were diluted in blocking buffer (1% Donkey Serum) and incubated overnight at 4°C. After three 5 min washes with PBS/ 0.1% Triton, secondary fluorescently labeled antibodies were diluted in blocking buffer (1% Donkey Serum) and incubated for 1 h at RT. Coverslips were washed two times with 1x PBS and an additional 2x with dH2O and mounted with ProLong® Gold antifade reagent (ThermoFisher).

### SDS PAGE and Western Blotting

Proteins were resolved on 4–20% Criterion TGX pre-cast gels (Biorad) and transferred to membranes by semi-dry trans-Blot Turbo transfer system (Biorad). The membranes were blocked with Odyssey Blocking Buffer (Licor Cat #927-40000) and then incubated for 1 h at RT or overnight at 4°C with the indicated primary antibody. The membranes were washed in TBST (3×5 min) followed by incubation for 1 h at RT with fluorescently conjugated goat anti-mouse, rat or rabbit IR Dye 680 or 800 antibodies (LICOR). The blots were washed in TBST (3×5 min) and scanned on an ODYSSEY^®^ CLx (LICOR). Quantitation of western blots was performed using Image Studio (LICOR).

### Sample preparation for FIB-SEM

Following airyscan microscopy, the cells were fixed and processed as previously described (*56*) with some differences. After dehydration the MatTek glass coverslip was removed from the plastic by using propylene oxide. The removed glass coverslip was then rinsed in 100% ethanol followed by immersion in mixtures of Durcupan ACM and ethanol with the following ratios: 25/75 for 1.5 h; 50/50 for 1.5 h; 75/25 overnight. The sample was then immersed in 100% Durcupan ACM for 4-5 hours with replacement of fresh Durcupan every hour. The glass coverslip was then removed and excess Durcupan was removed using filter paper. The coverslip was then placed in an oven at 60°C for 10 min, after which the sample was placed vertically in a 50 mL Falcon tube in folded filter papers and centrifuged for 15 min at 37°C and 750 RCF. The glass coverslip was then placed in an oven at 60°C under vacuum and left to polymerize over 2 days. The sample was then sputter coated with 50 nm gold and painted with silver paint, followed by drying under vacuum. The samples were imaged inside a Zeiss Crossbeam 540 FIB-SEM microscope. Platinum and carbon were deposited over the region of interest and the run was setup and controlled by Atlas software (Fibics). SEM settings: 1.5 kV; 2.0 nA; Milling probe: 300 pA. The slice thickness and voxel size were set to 6 nm. The total volume acquired (XYZ): 44.21 x 4.22 x 27.38 µm.

### Statistical analysis

Analyses based on cell counts were performed by an investigator blinded to treatment/transfection status. Statistical analysis for experiments with two treatment groups used Student’s t-tests with Welsh’s correction for unequal variance. For more than two groups, we used one-way ANOVA or two-way ANOVA where there were two factors in the model. Tukey’s *post-hoc* test was used to determine statistical significance for individual comparisons in those cases where the underlying ANOVA was statistically significant and where all groups were compared; Dunnett’s multiple comparison test was used where all groups were compared back to a single control group. Unless otherwise stated, graphed data are presented as means ± SEM. Comparisons were considered statistically significant where *p*< 0.05. *, *p*< 0.05; **, *p*< 0.01; ***, *p*< 0.001; ****, *p*< 0.0001.

## REFERENCES

1. C. Paisán-Ruíz, S. Jain, E. W. Evans, W. P. Gilks, J. Simón, M. van der Brug, A. López de Munain, S. Aparicio, A. M. Gil, N. Khan, J. Johnson, J. R. Martinez, D. Nicholl, I. Martí Carrera, A. S. Pena, R. de Silva, A. Lees, J. F. Martí-Massó, J. Pérez-Tur, N. W. Wood, A. B. Singleton, Cloning of the gene containing mutations that cause PARK8-linked Parkinson’s disease. Neuron. 44, 595–600 (2004).

2. S. Lesage, A.-L. Leutenegger, P. Ibanez, S. Janin, E. Lohmann, A. Dürr, A. Brice, French Parkinson’s Disease Genetics Study Group, LRRK2 haplotype analyses in European and North African families with Parkinson disease: a common founder for the G2019S mutation dating from the 13th century. Am. J. Hum. Genet. 77, 330–332 (2005).

3. J. Simón-Sánchez, C. Schulte, J. M. Bras, M. Sharma, J. R. Gibbs, D. Berg, C. Paisan-Ruiz, P. Lichtner, S. W. Scholz, D. G. Hernandez, R. Krüger, M. Federoff, C. Klein, A. Goate, J. Perlmutter, M. Bonin, M. A. Nalls, T. Illig, C. Gieger, H. Houlden, M. Steffens, M. S. Okun, B. A. Racette, M. R. Cookson, K. D. Foote, H. H. Fernandez, B. J. Traynor, S. Schreiber, S. Arepalli, R. Zonozi, K. Gwinn, M. van der Brug, G. Lopez, S. J. Chanock, A. Schatzkin, Y. Park, A. Hollenbeck, J. Gao, X. Huang, N. W. Wood, D. Lorenz, G. Deuschl, H. Chen, O. Riess, J. A. Hardy, A. B. Singleton, T. Gasser, Genome-wide association study reveals genetic risk underlying Parkinson’s disease. Nat. Genet. 41, 1308–1312 (2009).

4. D. Chang, M. A. Nalls, I. B. Hallgrímsdóttir, J. Hunkapiller, M. van der Brug, F. Cai, International Parkinson’s Disease Genomics Consortium, 23andMe Research Team, G. A. Kerchner, G. Ayalon, B. Bingol, M. Sheng, D. Hinds, T. W. Behrens, A. B. Singleton, T. R. Bhangale, R. R. Graham, A meta-analysis of genome-wide association studies identifies 17 new Parkinson’s disease risk loci. Nat. Genet. 49, 1511–1516 (2017).

5. A. M. A. Nalls, C. Blauwendraat, C. L. Vallerga, K. Heilbron, S. Bandres-Ciga, D. Chang, M. Tan, D. A. Kia, A. J. Noyce, A. Xue, J. Bras, E. Young, R. von Coelln, J. Simón-Sánchez, C. Schulte, M. Sharma, L. Krohn, L. Pihlstrøm, A. Siitonen, H. Iwaki, H. Leonard, F. Faghri, J. R. Gibbs, D. G. Hernandez, S. W. Scholz, J. A. Botia, M. Martinez, J.-C. Corvol, S. Lesage, J. Jankovic, L. M. Shulman, M. Sutherland, P. Tienari, K. Majamaa, M. Toft, O. A. Andreassen, T. Bangale, A. Brice, J. Yang, Z. Gan-Or, T. Gasser, P. Heutink, J. M. Shulman, N. W. Wood, D. Hinds, J. A. Hardy, H. R. Morris, J. Gratten, P. M. Visscher, R. R. Graham, A. B. Singleton, 23andMe Research Team, System Genomics of Parkinson’s Disease Consortium, International Parkinson’s Disease Genomics Consortium, Identification of novel risk loci, causal insights, and heritable risk for Parkinson’s disease: a meta-analysis of genome-wide association studies. Lancet Neurol. 18, 1091–1102 (2019).

6. E. Greggio, S. Jain, A. Kingsbury, R. Bandopadhyay, P. Lewis, A. Kaganovich, M. P. van der Brug, A. Beilina, J. Blackinton, K. J. Thomas, R. Ahmad, D. W. Miller, S. Kesavapany, A. Singleton, A. Lees, R. J. Harvey, K. Harvey, M. R. Cookson, Kinase activity is required for the toxic effects of mutant LRRK2/dardarin. Neurobiol. Dis. 23, 329–341 (2006).

7. A. B. West, D. J. Moore, S. Biskup, A. Bugayenko, W. W. Smith, C. A. Ross, V. L. Dawson, T. M. Dawson, Parkinson’s disease-associated mutations in leucine-rich repeat kinase 2 augment kinase activity. Proc. Natl. Acad. Sci. U. S. A. 102, 16842–16847 (2005).

8. A. Beilina, I. N. Rudenko, A. Kaganovich, L. Civiero, H. Chau, S. K. Kalia, L. V. Kalia, E. Lobbestael, R. Chia, K. Ndukwe, J. Ding, M. A. Nalls, International Parkinson’s Disease Genomics Consortium, North American Brain Expression Consortium, M. Olszewski, D. N. Hauser, R. Kumaran, A. M. Lozano, V. Baekelandt, L. E. Greene, J.-M. Taymans, E. Greggio, M. R. Cookson, Unbiased screen for interactors of leucine-rich repeat kinase 2 supports a common pathway for sporadic and familial Parkinson disease. Proc. Natl. Acad. Sci. U. S. A. 111, 2626–2631 (2014).

9. M. Steger, F. Tonelli, G. Ito, P. Davies, M. Trost, M. Vetter, S. Wachter, E. Lorentzen, G. Duddy, S. Wilson, M. A. Baptista, B. K. Fiske, M. J. Fell, J. A. Morrow, A. D. Reith, D. R. Alessi, M. Mann, Phosphoproteomics reveals that Parkinson’s disease kinase LRRK2 regulates a subset of Rab GTPases. Elife. 5 (2016), doi:10.7554/eLife.12813.

10. M. Steger, F. Diez, H. S. Dhekne, P. Lis, R. S. Nirujogi, O. Karayel, F. Tonelli, T. N. Martinez, E. Lorentzen, S. R. Pfeffer, D. R. Alessi, M. Mann, Systematic proteomic analysis of LRRK2-mediated Rab GTPase phosphorylation establishes a connection to ciliogenesis. Elife. 6 (2017), doi:10.7554/eLife.31012.

11. A. Ramirez, A. Heimbach, J. Gründemann, B. Stiller, D. Hampshire, L. P. Cid, I. Goebel, A. F. Mubaidin, A.-L. Wriekat, J. Roeper, A. Al-Din, A. M. Hillmer, M. Karsak, B. Liss, C. G. Woods, M. I. Behrens, C. Kubisch, Hereditary parkinsonism with dementia is caused by mutations in ATP13A2, encoding a lysosomal type 5 P-type ATPase. Nat. Genet. 38, 1184–1191 (2006).

12. A. D. Klein, J. R. Mazzulli, Is Parkinson’s disease a lysosomal disorder? Brain. 141, 2255–2262 (2018).

13. Y. Tong, H. Yamaguchi, E. Giaime, S. Boyle, R. Kopan, R. J. Kelleher 3rd, J. Shen, Loss of leucine-rich repeat kinase 2 causes impairment of protein degradation pathways, accumulation of alpha-synuclein, and apoptotic cell death in aged mice. Proc. Natl. Acad. Sci. U. S. A. 107, 9879–9884 (2010).

14. Y. Tong, E. Giaime, H. Yamaguchi, T. Ichimura, Y. Liu, H. Si, H. Cai, J. V. Bonventre, J. Shen, Loss of leucine-rich repeat kinase 2 causes age-dependent bi-phasic alterations of the autophagy pathway. Mol. Neurodegener. 7, 2 (2012).

15. L. Pellegrini, D. N. Hauser, Y. Li, A. Mamais, A. Beilina, R. Kumaran, A. Wetzel, J. Nixon-Abell, G. Heaton, I. Rudenko, M. Alkaslasi, N. Ivanina, H. L. Melrose, M. R. Cookson, K. Harvey, Proteomic analysis reveals co-ordinated alterations in protein synthesis and degradation pathways in LRRK2 knockout mice. Hum. Mol. Genet. 27, 3257–3271 (2018).

16. A. G. Henry, S. Aghamohammadzadeh, H. Samaroo, Y. Chen, K. Mou, E. Needle, W. D. Hirst, Pathogenic LRRK2 mutations, through increased kinase activity, produce enlarged lysosomes with reduced degradative capacity and increase ATP13A2 expression. Hum. Mol. Genet. 24, 6013–6028 (2015).

17. L. N. Hockey, B. S. Kilpatrick, E. R. Eden, Y. Lin-Moshier, G. C. Brailoiu, E. Brailoiu, C. E. Futter, A. H. Schapira, J. S. Marchant, S. Patel, Dysregulation of lysosomal morphology by pathogenic LRRK2 is corrected by TPC2 inhibition. J. Cell Sci. 128, 232–238 (2015).

18. S. J. Orenstein, S.-H. Kuo, I. Tasset, E. Arias, H. Koga, I. Fernandez-Carasa, E. Cortes, L. S. Honig, W. Dauer, A. Consiglio, A. Raya, D. Sulzer, A. M. Cuervo, Interplay of LRRK2 with chaperone-mediated autophagy. Nat. Neurosci. 16, 394– 406 (2013).

19. R. Wallings, N. Connor-Robson, R. Wade-Martins, LRRK2 interacts with the vacuolar-type H+-ATPase pump a1 subunit to regulate lysosomal function. Hum. Mol. Genet. 28, 2696–2710 (2019).

20. L. F. Burbulla, S. Jeon, J. Zheng, P. Song, R. B. Silverman, D. Krainc, A modulator of wild-type glucocerebrosidase improves pathogenic phenotypes in dopaminergic neuronal models of Parkinson’s disease. Sci. Transl. Med. 11 (2019), doi:10.1126/scitranslmed.aau6870.

21. R. C. Gomez, P. Wawro, P. Lis, D. R. Alessi, S. R. Pfeffer, Membrane association but not identity is required for LRRK2 activation and phosphorylation of Rab GTPases. J. Cell Biol. (2019), doi:10.1083/jcb.201902184.

22. T. Eguchi, T. Kuwahara, M. Sakurai, T. Komori, T. Fujimoto, G. Ito, S.-I. Yoshimura, A. Harada, M. Fukuda, M. Koike, T. Iwatsubo, LRRK2 and its substrate Rab GTPases are sequentially targeted onto stressed lysosomes and maintain their homeostasis. Proc. Natl. Acad. Sci. U. S. A. 115, E9115–E9124 (2018).

23. A. di Domenico, G. Carola, C. Calatayud, M. Pons-Espinal, J. P. Muñoz, Y. Richaud-Patin, I. Fernandez-Carasa, M. Gut, A. Faella, J. Parameswaran, J. Soriano, I. Ferrer, E. Tolosa, A. Zorzano, A. M. Cuervo, A. Raya, A. Consiglio, Patient-Specific iPSC-Derived Astrocytes Contribute to Non-Cell-Autonomous Neurodegeneration in Parkinson’s Disease. Stem Cell Reports. 12, 213–229 (2019).

24. K. Stafa, A. Trancikova, P. J. Webber, L. Glauser, A. B. West, D. J. Moore, GTPase Activity and Neuronal Toxicity of Parkinson’s Disease–Associated LRRK2 Is Regulated by ArfGAP1. PLoS Genetics. 8 (2012), p. e1002526.

25. A. Mamais, R. Chia, A. Beilina, D. N. Hauser, C. Hall, P. A. Lewis, M. R. Cookson, R. Bandopadhyay, Arsenite stress down-regulates phosphorylation and 14-3-3 binding of leucine-rich repeat kinase 2 (LRRK2), promoting self-association and cellular redistribution. J. Biol. Chem. 289, 21386–21400 (2014).

26. X.-T. Cheng, Y.-X. Xie, B. Zhou, N. Huang, T. Farfel-Becker, Z.-H. Sheng, Characterization of LAMP1-labeled nondegradative lysosomal and endocytic compartments in neurons. J. Cell Biol. 217, 3127–3139 (2018).

27. V. Hung, N. D. Udeshi, S. S. Lam, K. H. Loh, K. J. Cox, K. Pedram, S. A. Carr, A. Y. Ting, Spatially resolved proteomic mapping in living cells with the engineered peroxidase APEX2. Nat. Protoc. 11, 456–475 (2016).

28. C. H. Hsu, D. Chan, B. Wolozin, LRRK2 and the stress response: interaction with MKKs and JNK-interacting proteins. Neurodegener. Dis. 7, 68–75 (2010).

29. E.-J. Bae, D.-K. Kim, C. Kim, M. Mante, A. Adame, E. Rockenstein, A. Ulusoy, M. Klinkenberg, G. R. Jeong, J. R. Bae, C. Lee, H.-J. Lee, B.-D. Lee, D. A. Di Monte, E. Masliah, S.-J. Lee, LRRK2 kinase regulates α-synuclein propagation via RAB35 phosphorylation. Nat. Commun. 9, 3465 (2018).

30. G. Montagnac, J.-B. Sibarita, S. Loubéry, L. Daviet, M. Romao, G. Raposo, P. Chavrier, ARF6 Interacts with JIP4 to control a motor switch mechanism regulating endosome traffic in cytokinesis. Curr. Biol. 19, 184–195 (2009).

31. T. Tanaka, M. Iino, K. Goto, Knockdown of Sec8 enhances the binding affinity of c-Jun N-terminal kinase (JNK)-interacting protein 4 for mitogen-activated protein kinase kinase 4 (MKK4) and suppresses the phosphorylation of MKK4, p38, and JNK, thereby inhibiting apoptosis. FEBS J. 281, 5237–5250 (2014).

32. S. Aits, J. Kricker, B. Liu, A.-M. Ellegaard, S. Hämälistö, S. Tvingsholm, E. Corcelle-Termeau, S. Høgh, T. Farkas, A. Holm Jonassen, I. Gromova, M. Mortensen, M. Jäättelä, Sensitive detection of lysosomal membrane permeabilization by lysosomal galectin puncta assay. Autophagy. 11, 1408–1424 (2015).

33. D. L. Thiele, P. E. Lipsky, Mechanism of L-leucyl-L-leucine methyl ester-mediated killing of cytotoxic lymphocytes: dependence on a lysosomal thiol protease, dipeptidyl peptidase I, that is enriched in these cells. Proc. Natl. Acad. Sci. U. S. A. 87, 83–87 (1990).

34. I. Maejima, A. Takahashi, H. Omori, T. Kimura, Y. Takabatake, T. Saitoh, A. Yamamoto, M. Hamasaki, T. Noda, Y. Isaka, T. Yoshimori, Autophagy sequesters damaged lysosomes to control lysosomal biogenesis and kidney injury. EMBO J. 32, 2336–2347 (2013).

35. H. Kobayashi, K. Etoh, N. Ohbayashi, M. Fukuda, Rab35 promotes the recruitment of Rab8, Rab13 and Rab36 to recycling endosomes through MICAL-L1 during neurite outgrowth. Biol. Open. 3, 803–814 (2014).

36. V. Marchesin, A. Castro-Castro, C. Lodillinsky, A. Castagnino, J. Cyrta, H. Bonsang-Kitzis, L. Fuhrmann, M. Irondelle, E. Infante, G. Montagnac, F. Reyal, A. Vincent-Salomon, P. Chavrier, ARF6-JIP3/4 regulate endosomal tubules for MT1-MMP exocytosis in cancer invasion. J. Cell Biol. 211, 339–358 (2015).

37. A. Zimprich, S. Biskup, P. Leitner, P. Lichtner, M. Farrer, S. Lincoln, J. Kachergus, M. Hulihan, R. J. Uitti, D. B. Calne, A. J. Stoessl, R. F. Pfeiffer, N. Patenge, I. C. Carbajal, P. Vieregge, F. Asmus, B. Müller-Myhsok, D. W. Dickson, T. Meitinger, T. M. Strom, Z. K. Wszolek, T. Gasser, Mutations in LRRK2 cause autosomal-dominant parkinsonism with pleomorphic pathology. Neuron. 44, 601–607 (2004).

38. J. Alegre-Abarrategui, H. Christian, M. M. P. Lufino, R. Mutihac, L. L. Venda, O. Ansorge, R. Wade-Martins, LRRK2 regulates autophagic activity and localizes to specific membrane microdomains in a novel human genomic reporter cellular model. Human Molecular Genetics. 18 (2009), pp. 4022–4034.

39. S. Biskup, D. J. Moore, F. Celsi, S. Higashi, A. B. West, S. A. Andrabi, K. Kurkinen, S.-W. Yu, J. M. Savitt, H. J. Waldvogel, R. L. M. Faull, P. C. Emson, R. Torp, O. P. Ottersen, T. M. Dawson, V. L. Dawson, Localization of LRRK2 to membranous and vesicular structures in mammalian brain. Ann. Neurol. 60, 557–569 (2006).

40. Z. Berger, K. A. Smith, M. J. Lavoie, Membrane localization of LRRK2 is associated with increased formation of the highly active LRRK2 dimer and changes in its phosphorylation. Biochemistry. 49, 5511–5523 (2010).

41. D. A. MacLeod, H. Rhinn, T. Kuwahara, A. Zolin, G. Di Paolo, B. D. McCabe, K. S. Marder, L. S. Honig, L. N. Clark, S. A. Small, A. Abeliovich, RAB7L1 interacts with LRRK2 to modify intraneuronal protein sorting and Parkinson’s disease risk. Neuron. 77, 425–439 (2013).

42. L. Bonet-Ponce, M. R. Cookson, The role of Rab GTPases in the pathobiology of Parkinson’ disease. Curr. Opin. Cell Biol. 59, 73–80 (2019).

43. E. Purlyte, H. S. Dhekne, A. R. Sarhan, R. Gomez, P. Lis, M. Wightman, T. N. Martinez, F. Tonelli, S. R. Pfeffer, D. R. Alessi, Rab29 activation of the Parkinson’s disease-associated LRRK2 kinase. EMBO J. 38 (2019), doi:10.15252/embj.2018101237.

44. K. E. McNally, P. J. Cullen, Endosomal Retrieval of Cargo: Retromer Is Not Alone. Trends Cell Biol. 28, 807–822 (2018).

45. S. Sridhar, B. Patel, D. Aphkhazava, F. Macian, L. Santambrogio, D. Shields, A. M. Cuervo, The lipid kinase PI4KIIIβ preserves lysosomal identity. EMBO J. 32, 324– 339 (2013).

46. L. Yu, C. K. McPhee, L. Zheng, G. A. Mardones, Y. Rong, J. Peng, N. Mi, Y. Zhao, Z. Liu, F. Wan, D. W. Hailey, V. Oorschot, J. Klumperman, E. H. Baehrecke, M. J. Lenardo, Termination of autophagy and reformation of lysosomes regulated by mTOR. Nature. 465, 942–946 (2010).

47. S. Pickles, P. Vigié, R. J. Youle, Mitophagy and Quality Control Mechanisms in Mitochondrial Maintenance. Curr. Biol. 28, R170–R185 (2018).

48. G.-L. McLelland, V. Soubannier, C. X. Chen, H. M. McBride, E. A. Fon, Parkin and PINK1 function in a vesicular trafficking pathway regulating mitochondrial quality control. EMBO J. 33, 282–295 (2014).

49. G.-L. McLelland, S. A. Lee, H. M. McBride, E. A. Fon, Syntaxin-17 delivers PINK1/parkin-dependent mitochondrial vesicles to the endolysosomal system. J. Cell Biol. 214, 275–291 (2016).

50. E. Greggio, I. Zambrano, A. Kaganovich, A. Beilina, J.-M. Taymans, V. Daniëls, P. Lewis, S. Jain, J. Ding, A. Syed, K. J. Thomas, V. Baekelandt, M. R. Cookson, The Parkinson disease-associated leucine-rich repeat kinase 2 (LRRK2) is a dimer that undergoes intramolecular autophosphorylation. J. Biol. Chem. 283, 16906–16914 (2008).

51. B. Huang, A. Hubber, J. A. McDonough, C. R. Roy, M. A. Scidmore, J. A. Carlyon, The Anaplasma phagocytophilum-occupied vacuole selectively recruits Rab-GTPases that are predominantly associated with recycling endosomes. Cell. Microbiol. 12, 1292–1307 (2010).

52. K. A. Rzomp, L. D. Scholtes, B. J. Briggs, G. R. Whittaker, M. A. Scidmore, Rab GTPases are recruited to chlamydial inclusions in both a species-dependent and species-independent manner. Infect. Immun. 71, 5855–5870 (2003).

53. A. S. S. Lam, J. D. Martell, K. J. Kamer, T. J. Deerinck, M. H. Ellisman, V. K. Mootha, Y. Ting, Directed evolution of APEX2 for electron microscopy and proximity labeling. Nat. Methods. 12, 51–54 (2015).

54. N. M. Sherer, M. J. Lehmann, L. F. Jimenez-Soto, A. Ingmundson, S. M. Horner, G. Cicchetti, P. G. Allen, M. Pypaert, J. M. Cunningham, W. Mothes, Visualization of retroviral replication in living cells reveals budding into multivesicular bodies. Traffic. 4, 785–801 (2003).

55. K. S. Conrad, T.-W. Cheng, D. Ysselstein, S. Heybrock, L. R. Hoth, B. A. Chrunyk, C. W. Am Ende, D. Krainc, M. Schwake, P. Saftig, S. Liu, X. Qiu, M. D. Ehlers, Lysosomal integral membrane protein-2 as a phospholipid receptor revealed by biophysical and cellular studies. Nat. Commun. 8, 1908 (2017).

56. C. K. E. Bleck, Y. Kim, T. Bradley Willingham, B. Glancy, Subcellular connectomic analyses of energy networks in striated muscle. Nature Communications. 9 (2018), doi:10.1038/s41467-018-07676-y.

