## Supplementary figures and images for "LRRK2 mediates tubulation and vesicle sorting from membrane damaged lysosomes"

### Suppl Figure 1

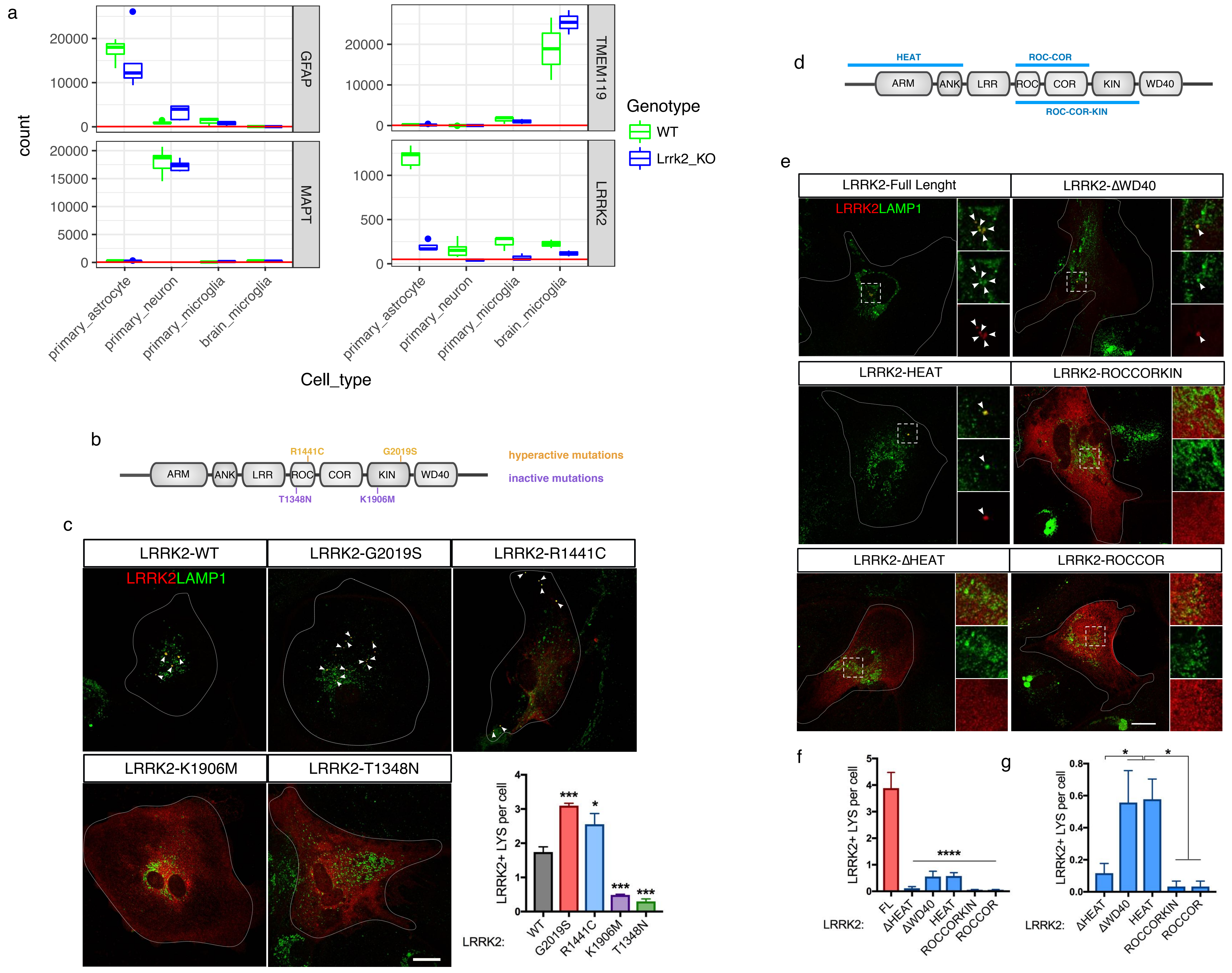


*Bonet-Ponce et al., Supp Figure 1*

### Suppl Figure 2

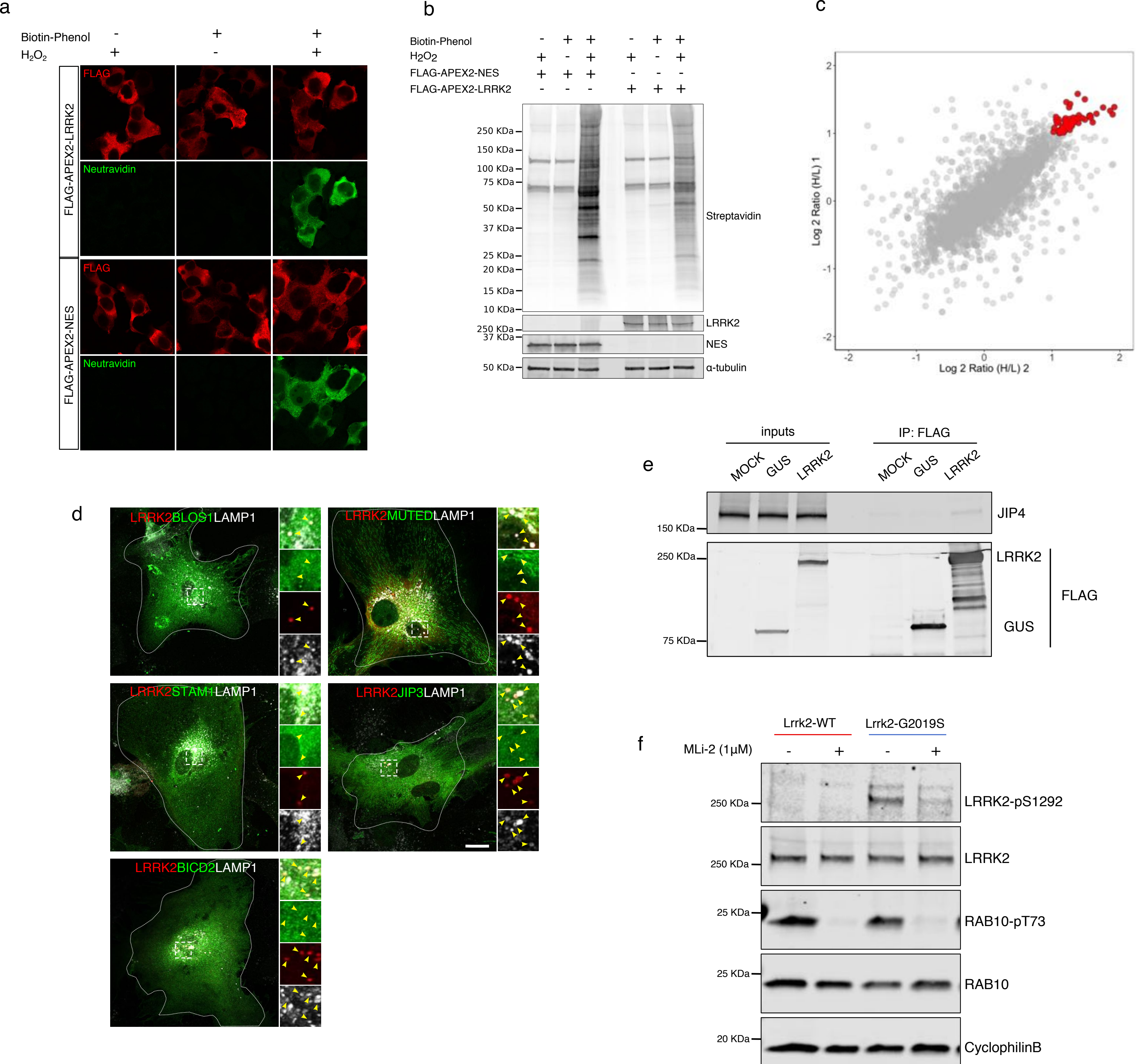


*Bonet-Ponce et al., Suppl Figure 2*

### Suppl Figure 3

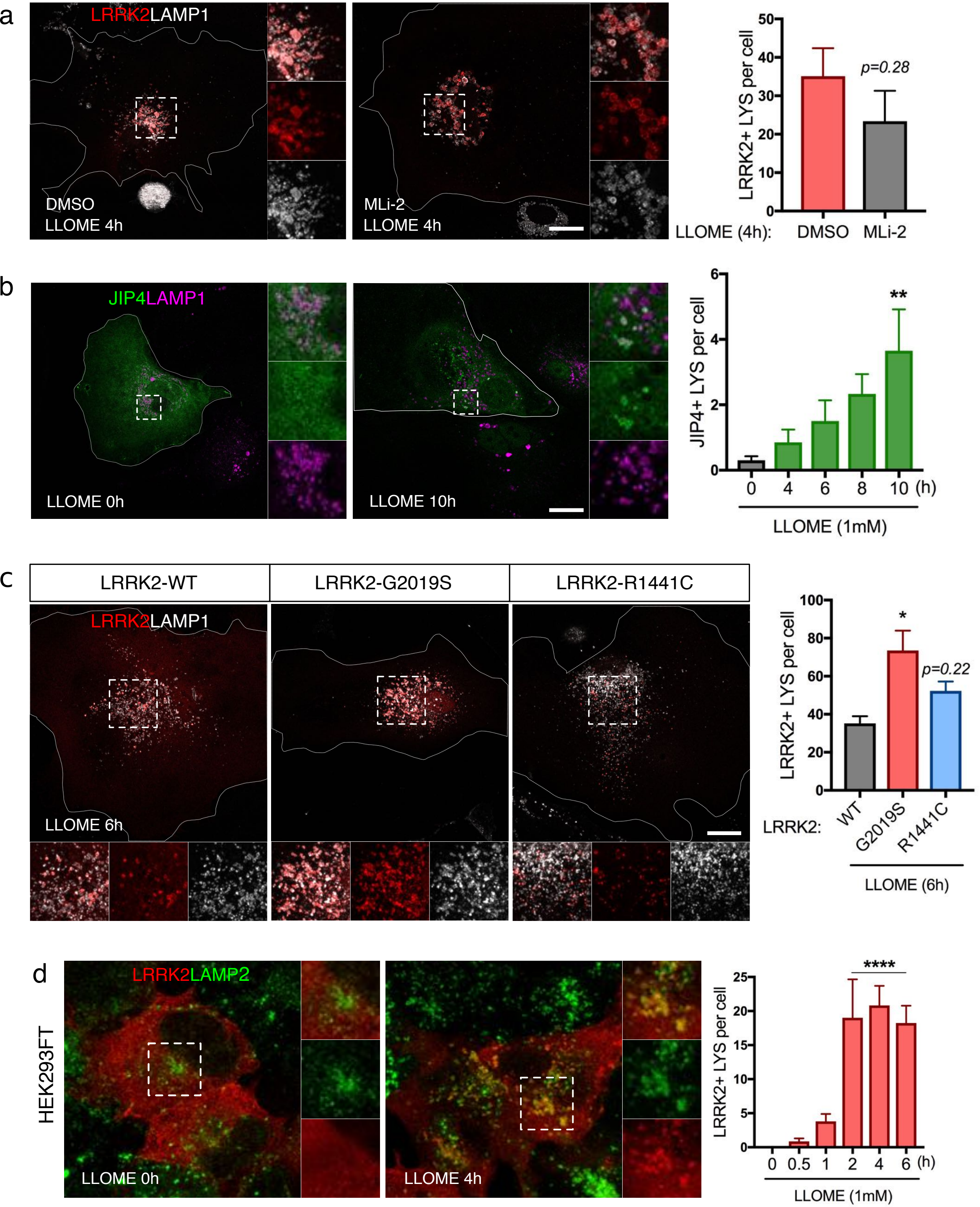


*Bonet-Ponce et al., Suppl Figure 3*

### Suppl Figure 4

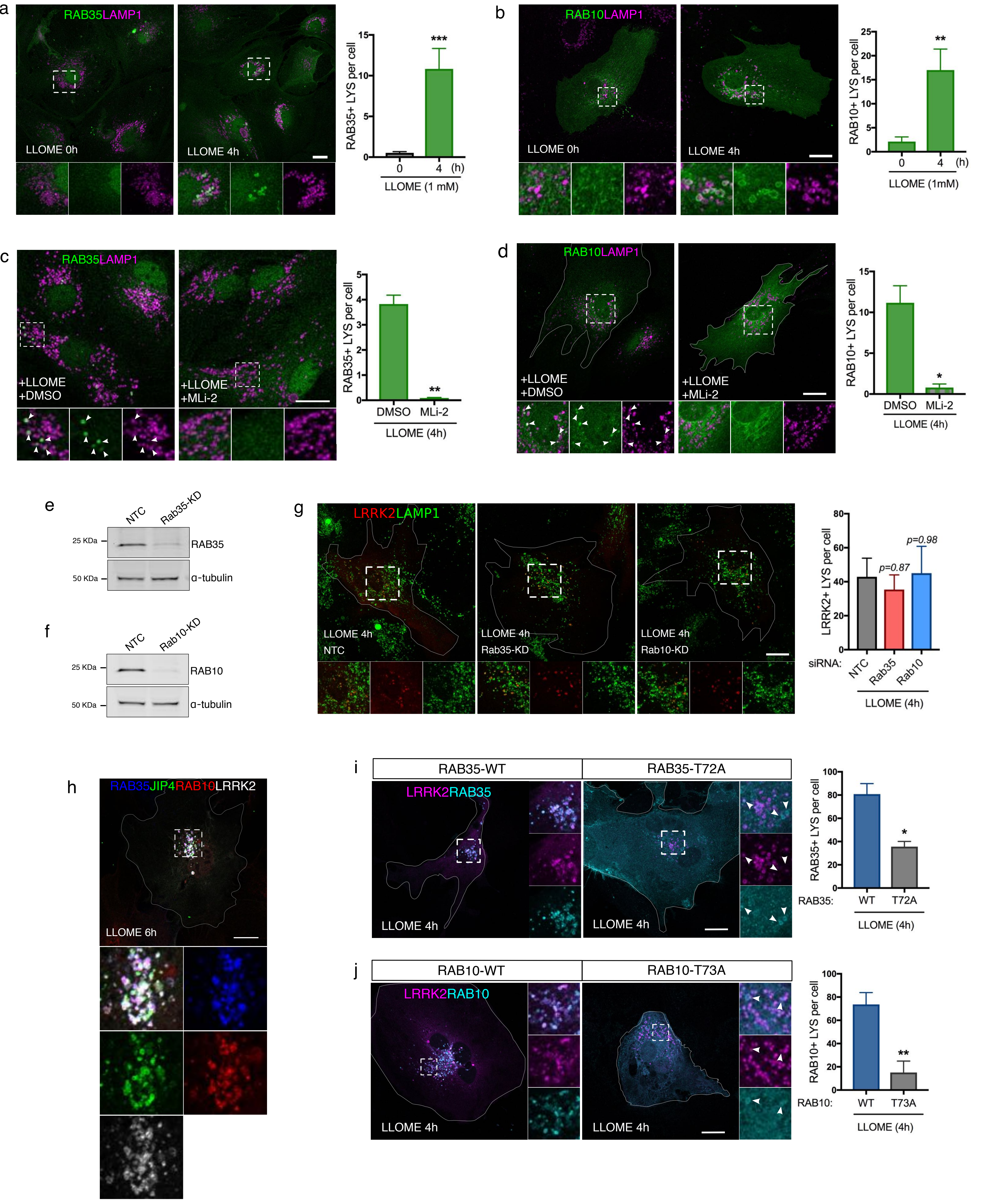


*Bonet-Ponce et al., Suppl Figure 4*

### Suppl Figure 5

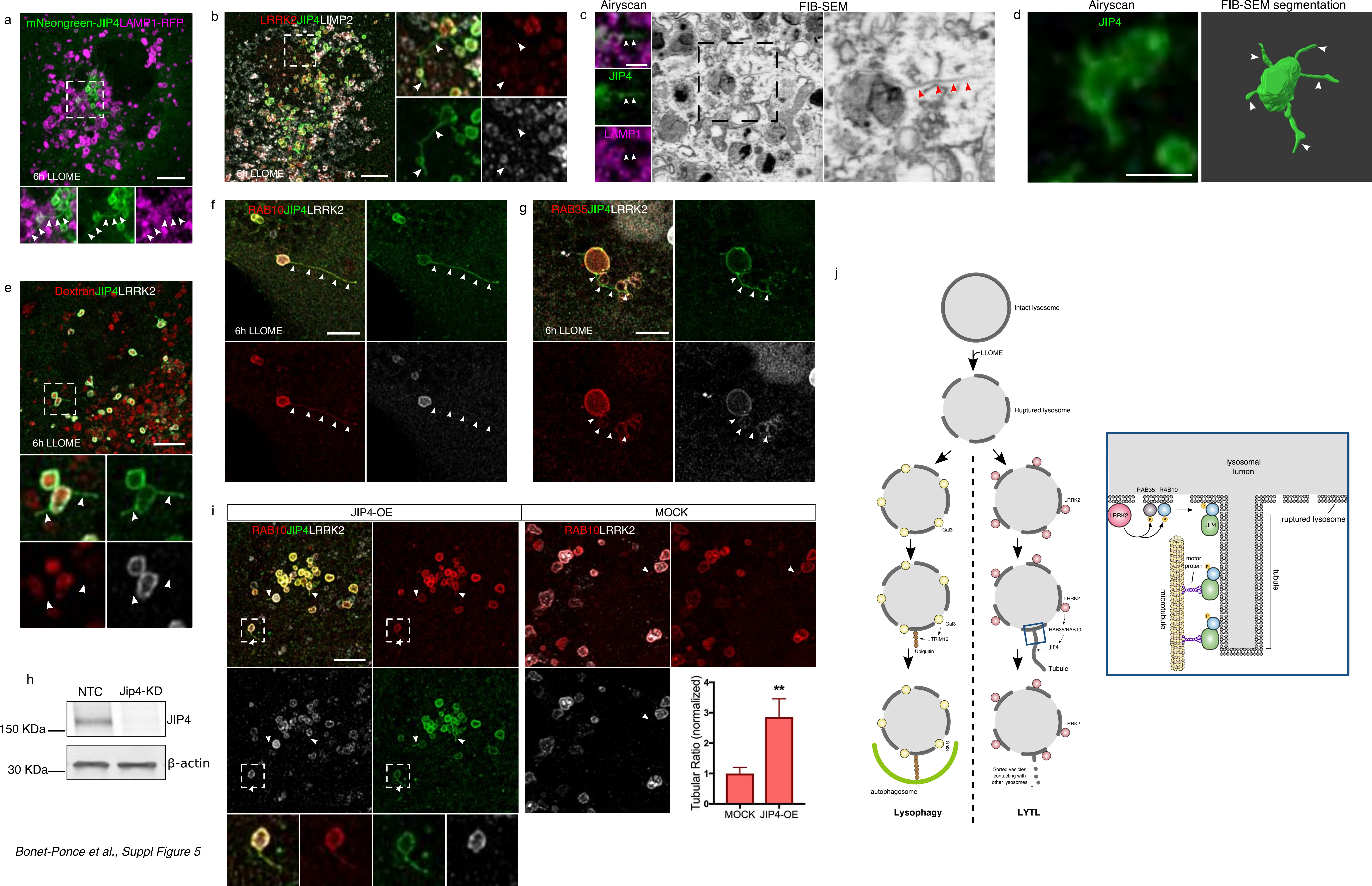


*Bonet-Ponce et al., Suppl Figure 5*
